# Regulation of transcription patterns, poly-ADP-ribose, and RNA-DNA hybrids by the ATM protein kinase

**DOI:** 10.1101/2023.12.06.570417

**Authors:** Phillip R. Woolley, Xuemei Wen, Olivia M. Conway, Nicolette A. Ender, Ji-Hoon Lee, Tanya T. Paull

## Abstract

The ATM protein kinase is a master regulator of the DNA damage response and also an important sensor of oxidative stress. Analysis of gene expression in Ataxia-telangiectasia patient brain tissue shows that large-scale transcriptional changes occur in patient cerebellum that correlate with expression level and GC content of transcribed genes. In human neuron-like cells in culture we map locations of poly-ADP-ribose and RNA-DNA hybrid accumulation genome-wide with ATM inhibition and find that these marks also coincide with high transcription levels, active transcription histone marks, and high GC content. Antioxidant treatment reverses the accumulation of R-loops in transcribed regions, consistent with the central role of ROS in promoting these lesions. Based on these results we postulate that transcription-associated lesions accumulate in ATM-deficient cells and that the single-strand breaks and PARylation at these sites ultimately generate changes in transcription that compromise cerebellum function and lead to neurodegeneration over time in A-T patients.

## Introduction

Ataxia-telangiectasia (A-T) is a childhood-onset progressive neurodegenerative disorder caused by the loss of the ATM protein kinase^1^. ATM is well-known as a cell cycle checkpoint kinase that responds to DNA damage in the form of double-strand breaks through the Mre11-Rad50-Nbs1 (MRN) complex and phosphorylates hundreds of substrates involved in checkpoints, DNA repair, and DNA damage survival^2,3^. In addition, we have previously shown that ATM can be activated independently of DNA breaks through direct oxidation of cysteine residues^4^ and many groups have noted the chronic oxidative stress in mammalian cells caused by ATM loss^3,5^. The molecular basis of ATM activation via oxidation has also been recently determined^6^. Despite the molecular insights into ATM function over the last two decades, however, it is still not clear how the loss of this kinase results in the cerebellum-specific neurodegeneration that is considered to be the most debilitating clinical outcome of this autosomal recessive disorder.

During our characterization of mutant ATM alleles in human cells and experiments with purified proteins in vitro, we identified several forms of ATM that fail to be activated by oxidative stress but have normal MRN-mediated activation^4,7^. One of these, the R3047X mutant lacking the last 10 amino acids of ATM, is responsible for cerebellar neurodegeneration in several patients^8–10^. For this reason we have focused our recent efforts on the ROS-dependent pathway of ATM activation, which we found in human tumor-derived cells to control many cellular responses including ROS levels, mitochondrial functions, ROS-induced checkpoints and autophagy, as well as protection from nuclear protein aggregate formation^11^. We also observed DNA damage in cells lacking ATM ROS activation, specifically single-strand DNA (ssDNA) breaks—a form of DNA damage known to be associated with cerebellar ataxia in several familial disorders^12,13^. The ssDNA breaks we observed with the loss of ATM activity or the loss of its ROS activation are transcription-dependent and are eliminated with overexpression of either RNaseH or the human Senataxin enzyme, both of which act to remove RNA-DNA hybrids^14^. The ssDNA breaks also induce hyper-poly-ADP-ribosylation (hyperPARylation) by PARP1 and PARP2 in ATM-deficient cells that drives the protein aggregation we observed^11^.

Based on these findings, we hypothesize that high ROS in human cells lacking ATM function causes transcriptional stress in the form of RNA-DNA hybrids that ultimately lead to ssDNA breaks and hyperPARylation. We further postulate that transcription patterns at specific genomic locations may be altered in A-T tissues because of the accumulated hybrids and DNA damage over time. To test these ideas directly we quantified transcripts in A-T patient cerebellum tissues using RNAseq and found significant changes between the patient and control groups that were primarily evident in the cerebellum. Analysis of these changes indicates a strong relationship between DNA sequence context, expression levels, and dysregulation in A-T patients. In addition, we used human neuron-like cells differentiated in culture to assess the locations of DNA damage in the absence of ATM function by quantifying levels of PAR and RNA-DNA hybrids genome-wide. Through this analysis we find patterns of DNA damage that increase with loss of ATM activity and correlate with both GC content and transcription levels. Taken together, these results suggest a unified model for deficiencies in critical gene expression patterns that occur as a result of ATM loss in human A-T patients.

## Results

### A-T cerebellum transcriptomics indicate significant changes in comparison with controls

Our previous results in human cells in culture and our analysis of A-T patient brain tissue by mass spectrometry^11^ suggested that transcription patterns may be altered in A-T patients. To quantify these changes we performed total RNA extraction and RNA sequencing on five A-T patient and four control patient brain tissues (Table S1). The tissue types examined included the cerebellum which is most affected functionally in A-T patients as well as two regions in the cortex which are not functionally affected (frontal cortex regions BA9 and BA10). Significant transcription differences were observed in the A-T patient cerebellum and BA9 compared to control cerebellum samples, whereas fewer changes were observed in A-T patient BA10 cortex samples (Figure 1A-C). We identified both upregulated genes (>2-fold compared to controls, q-value<0.01, 714 transcripts) and downregulated genes (<2-fold compared to controls, q-value<0.01, 1151 transcripts) in A-T patient cerebellum tissue (Figure 1D). A substantial subset of these differentially expressed genes were found to be more than 4-fold up or downregulated (486 and 477, respectively). Gene set enrichment analysis revealed many pathways affected in cerebellum and cortex tissues, including calcium signaling, oxidative phosphorylation, and pathways related to other forms of neurodegeneration (Figure S1).

**Figure 1.**
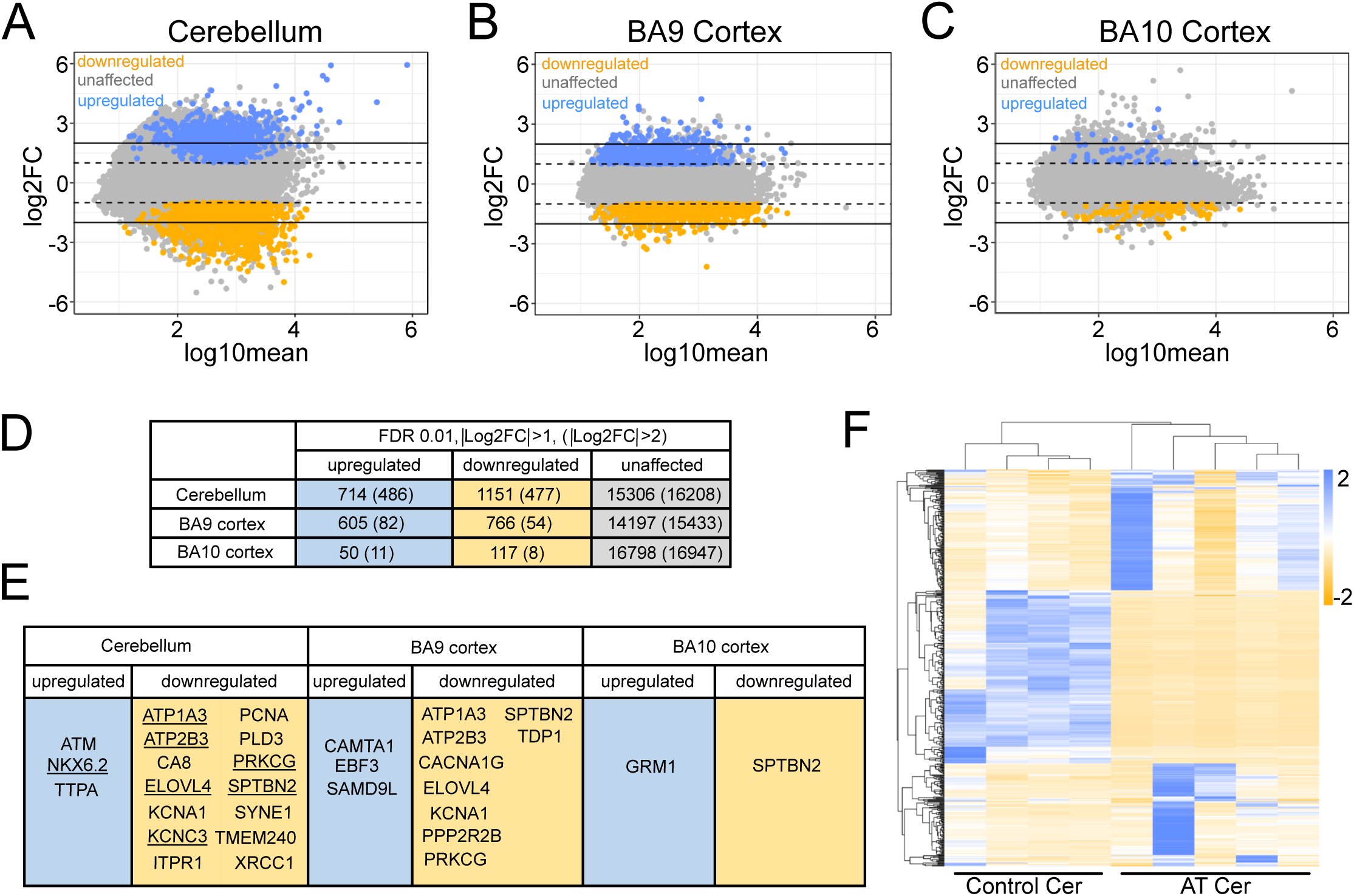
The A-T patient transcriptome shows dramatic differences from controls. (A) MA plot comparing cerebellar AT patient transcriptome to control patient transcriptome. Dashed lines correspond to |Log2FC|>1, solid lines correspond to |Log2FC|>2. Orange corresponds to Log2FC<-1 with qvalue<0.01 (downregulated gene set) and blue corresponds to Log2FC>1 with qvalue<0.01 (upregulated gene set). (B) MA plot comparing BA9 cortex AT patient transcriptome to control patient transcriptome. (C) MA plot comparing BA10 cortex AT patient transcriptome to control patient transcriptome. (D) Numbers of genes in upregulated (|Log2FC|>1), downregulated (Log2FC<-1), and unaffected gene sets in each tissue type. Gene sets in paratheses indicate numbers of differentially transcribed genes with at least 4-fold change from wild-type (|Log2FC|>2). (E) Human cerebellar ataxia-linked genes that fall within upregulated and downregulated gene sets for each tissue type. Underlined genes are in the |Log2FC|>2 subset. (F) Heatmap for the 500 genes with the most extreme |Log2FC| values in the cerebellum. Columns are patients, rows are genes.

Notably, transcripts with significant downregulation in the cerebellum and BA10 cortex included several genes linked to other forms of ataxia (Figure 1E). These include *ITPR1* and *CA8*, genes involved in in inositol phosphate regulation of calcium signaling that we previously found to be reduced at the protein level^11^. In addition to these, 12 other genes we identified in the subset of transcripts strongly reduced in A-T patients are individually linked to cerebellar ataxia in other hereditary disorders^15–29^.

In addition to the large number of dysregulated genes common to all the A-T patients tested, heatmap analysis also demonstrated significant heterogeneity between A-T patient cerebellum samples (Figure 1F). The level of transcriptional change specific to each patient were not obviously correlated with the age or sex of the patient.

Given the magnitude of transcriptional changes observed in the patients and the large number of genes affected, we sought to identify characteristics that distinguish affected from unaffected genes. We grouped transcripts according to their levels in A-T patients: downregulated, unaffected, or upregulated compared to controls. Comparisons of gene groups in relation to their gene expression level in controls showed an obvious separation between the downregulated and upregulated genes. Specifically, genes downregulated in A-T tended to be those that are highly expressed in control patients whereas genes upregulated in A-T tended to be lowly expressed in control patients (levels of transcript change in A-T patients relative to controls shown in relation to the rank of each transcript’s expression in control individuals, Figure 2A). Comparisons between these subsets passed non-parametric cumulative distribution tests including Kolmogorov-Smirnov and Anderson-Darling with 5000 bootstraps. The highly expressed genes, largely downregulated in patients, include many transcripts specific to Purkinje cells that are considered to be one of the most affected cell types in A-T^1^. Another way to view these data is to show the downregulated or upregulated transcripts in expression quantiles in comparison to transcript levels in normal cells. This analysis shows that the genes downregulated in A-T are biased toward those that are highly expressed in the cerebellum (Figure 2B), while the genes upregulated in A-T are biased toward those that are expressed at a low level (Figure 2C). Similar trends are observed in the cortex tissue of A-T patients, with the downregulated gene set being among the highest expressed in that tissue, although there are fewer genes affected and the trends are not as extreme as observed in the cerebellum (Figure 2D-F).

**Figure 2.**
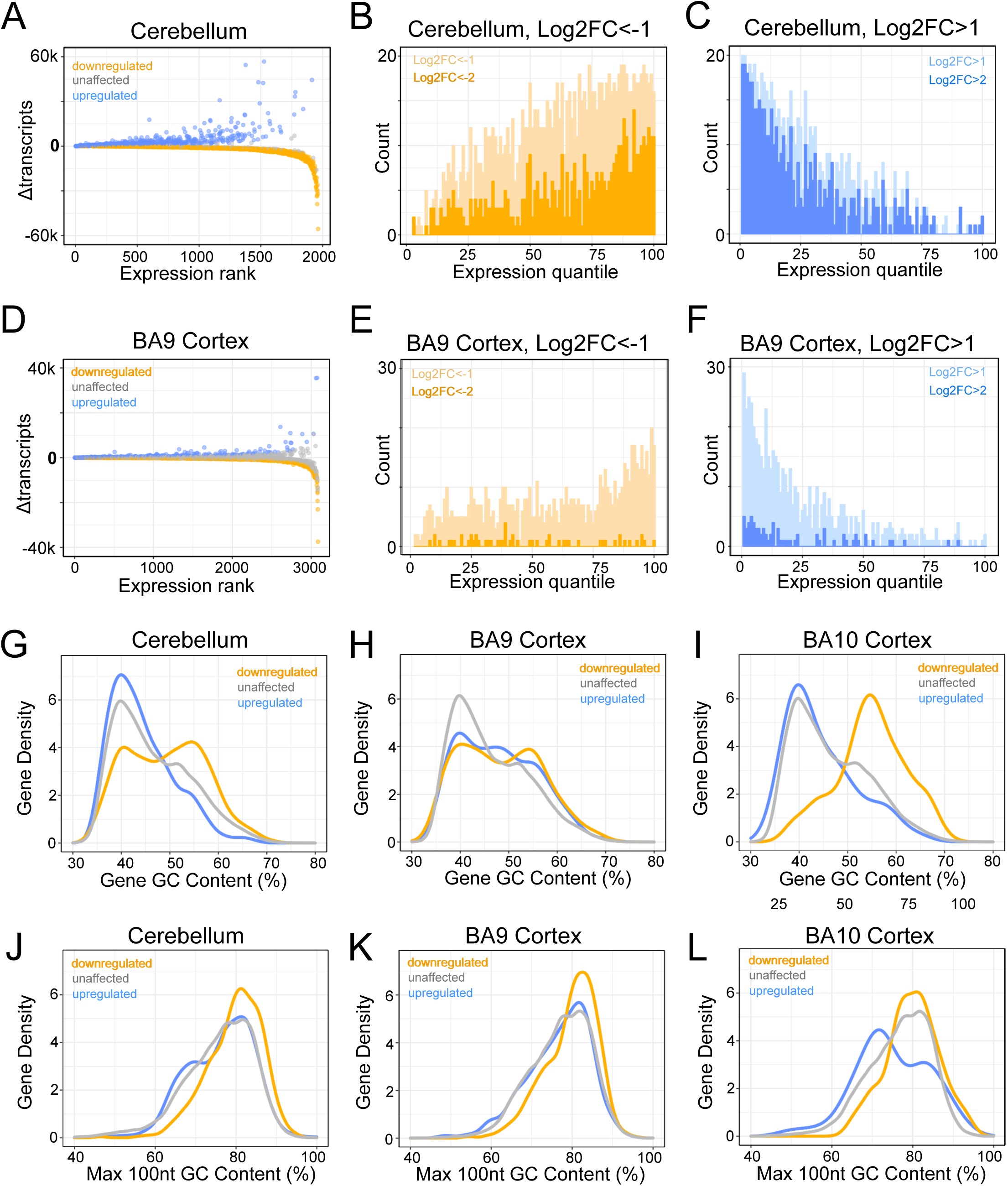
Differentially expressed genes in A-T patients align with patterns of transcript abundance and GC content. (A) Transcript change vs expression rank plots for cerebellum genes with qvalue<0.01. Expression rank is defined here as the set of all genes with q-value<0.01 sorted by mean control patient transcript abundance. The higher the expression rank, the higher expressed the gene is in control patients. (B) Expression quantile histograms for cerebellum downregulated gene set. As in (A) but without a q-value requirement, all genes were sorted by mean control patient transcript abundance before being divided into 100 equally sized, nonoverlapping bins. The rightmost bin contains the 1% highest expressed genes in control patients, the leftmost bin contains the 1% lowest expressed genes in control patients, etc. The number of downregulated genes in each bin are plotted on the histogram. Light orange: genes with Log2FC<1 in A-T patients relative to controls; dark orange: genes with Log2FC<2. (C) Expression quantile histograms for cerebellum upregulated gene set. Light blue: genes with Log2FC>1 in A-T patients relative to controls; dark blue: genes with Log2FC>2. (D). As in (A), but for the BA9 cortex. (E) As in (B), but for the BA9 cortex. (F) As in (C), but for the BA9 cortex. (G) Density plots for gene GC content for cerebellar upregulated, downregulated, and unaffected gene sets from Fig 1A. (H) Density plots for gene GC content for BA9 upregulated, downregulated, and unaffected gene sets from Fig 1B. (I) Density plots for gene GC content for BA10 upregulated, downregulated, and unaffected gene sets from Fig 1C. (J) Density plots for the maximum 100nt GC content per gene for cerebellar upregulated, downregulated, and unaffected gene sets. (K) Density plots for the maximum 100nt GC content per gene for BA9 upregulated, downregulated, and unaffected gene sets. (L) Density plots for the maximum 100nt GC content per gene for BA10 upregulated, downregulated, and unaffected gene sets.

Although the affected genes in A-T patient tissues are not all specific to Purkinje cells, it is possible that one explanation for these trends could be the reduced density of Purkinje cells in patient tissue. To investigate this we re-analyzed micrographs of A-T patient cerebellum that were collected in our previous study^11^(Table S1). 6 A-T patient tissue samples and 6 controls were analyzed for the density of Purkinje cells by examining previously collected images of the tissues and determining the number of cells per micron of Purkinje layer. These results showed that A-T patients exhibit a reduction of ∼40% in the density of Purkinje neurons in the Purkinje layer (Figure S2). This difference is significant based on these 6 pairs of patients and controls, although the loss of mRNA transcripts observed in patients is much more extreme (changes leading to loss of >75% shown in Figures 1 and 2). Notably, the expression level downregulation trend was also observed in the BA9 cortex tissue which does not have any precedent of neurodegeneration in A-T patients and does not show cellular atrophy^30^.

Lastly, we analyzed the GC content of each class of genes (upregulated, downregulated, or unchanged in A-T patients) and found that the downregulated genes show a distinct pattern of high overall GC content per gene (Figure 2G-I) and higher than average GC content when considering the 100nt of maximum GC content per gene (Figure 2J-L). In contrast, the unaffected and upregulated genes show lower GC content overall and lower than average GC content when considering the 100nt of maximum GC content per gene. These patterns are most obvious in the cerebellum tissue but also apparent in the neocortex samples. Comparisons of downregulated with non-downregulated gene sets passed non-parametric cumulative distribution tests including Kolmogorov-Smirnov and Anderson-Darling with 5000 bootstraps. In contrast, no differences were observed between differentially expressed genes with respect to GC skew (defined as *(G − C)/(G + C*)) regardless of whether the entire gene or just the promoter region was included in this analysis (Figure S3). Overall, the analysis suggests that the patterns of gene expression in A-T patient tissue are non-random and strongly correlated with both the level of gene expression in normal individuals as well as DNA sequence context.

To confirm and validate these findings using an orthogonal method, we used spatial transcriptomics to analyze a cerebellum tissue slice from an A-T patient in comparison to a control individual. These results show striking differences between the patient and control tissue, particularly in the molecular layer (Figure S4). Although Purkinje cells cannot be uniquely analyzed from this method due to low resolution, we observed downregulation of many genes known to be specific to this cell type, including ITPR1 and CA8.

### Quantification of PAR genome-wide in ATM-inhibited neuron-like cells

From our previous study of the effects of ATM on tumor-derived cells in culture we know that loss of ATM activity leads to high levels of reactive oxygen species (ROS) and that RNA-DNA hybrids accumulate as a result of high ROS^11^. In addition, we observed single-strand breaks and hyperactivation of poly-ADP-ribose polymerases 1 and 2 (PARP1 and 2) in these cells with loss of ATM or its oxidative activation pathway. Based on these findings we hypothesize that RNA-DNA hybrids and PARP activation occurs at sites of transcriptional stress in ATM-deficient cells and that these events likely lead to transcriptional changes over time as we observed in A-T patients (working model in Figure 3A).

**Figure 3.**
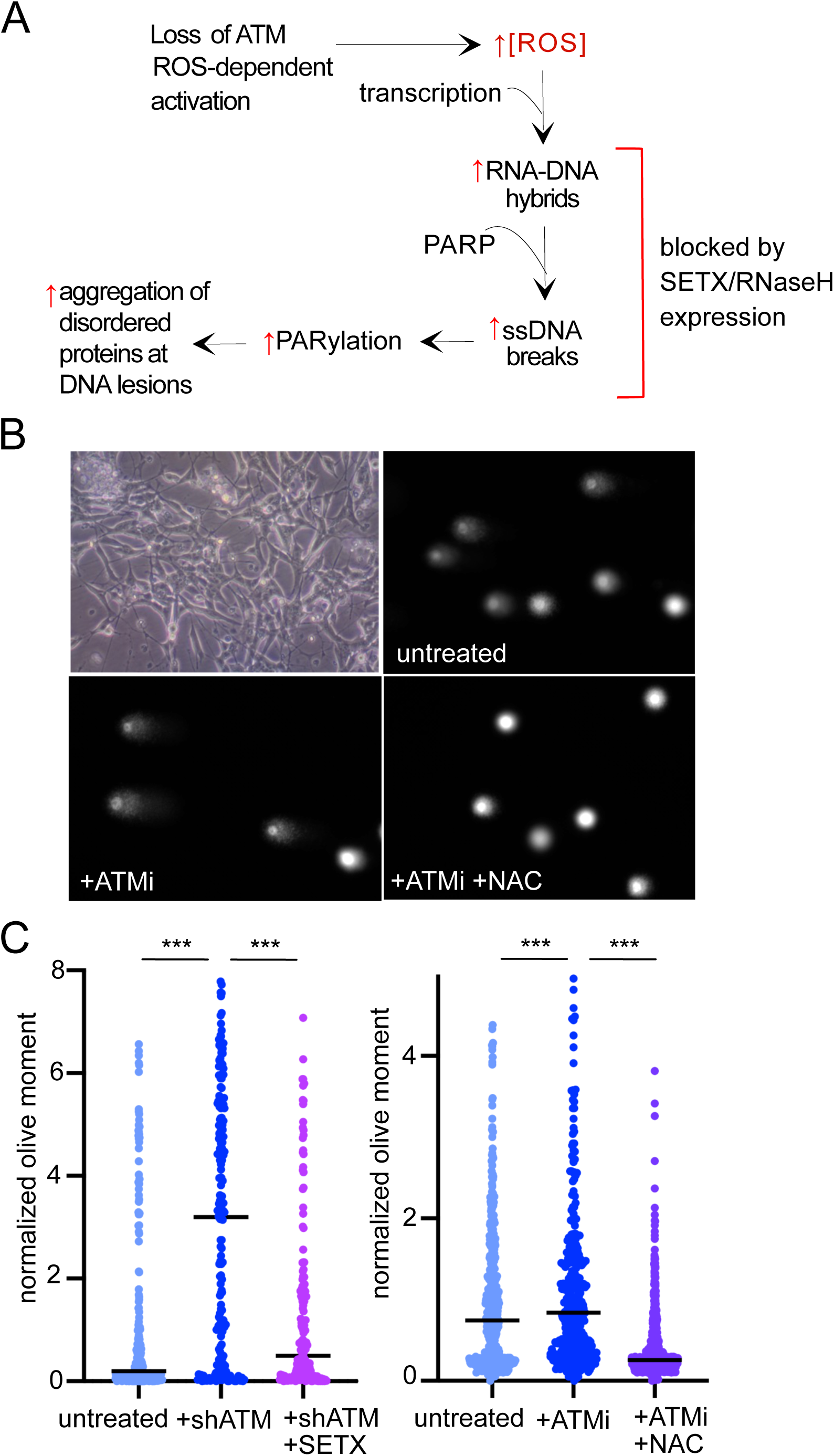
ATM inhibition in post-mitotic neuronal-like cells generates ROS-dependent single-strand breaks. (A) Working model for transcription-associated lesions in ATM-deficient cells based on previous observations^7,11^. Loss of ATM produces high ROS in human cells which generates R-loops, single-strand DNA breaks, hyperPARylation, and protein aggregates. (B) Brightfield image of differentiated cells (top left) and examples of alkaline comet assay results from post-mitotic human neuron-like cells differentiated from SH-SY5Y neuroblastoma cells comparing untreated, ATM inhibitor-treated (ATMi), and ATMi + N-acetyl cysteine (NAC) conditions. (C) Quantitation of the olive moment in alkaline comet assays measuring > 380 cells from each treatment group. Differentiated SH-SY5Y cells were depleted of ATM by shRNA and complemented with expression of the C-terminal helicase domain of SETX as indicated (left panel). Differentiated SH-SY5Y cells were treated with ATMi or ATMi with NAC as indicated (right panel). *** p<0.0005 by 2-sample t-test assuming unequal variances.

To test this hypothesis we employed human SH-SY5Y neuroblastoma cells that were differentiated into post-mitotic neuronal-like cells as previously described over the course of 7 days using retinoic acid and BDNF^11,31^. Under these conditions, inhibition of ATM in differentiated SH-SY5Y cells generates significantly higher levels of ROS (Figure S5) as well as single-strand DNA breaks as measured by alkaline comet assay (Figure 3B,C). No increase in double-strand breaks was observed under these conditions (Figure S6). As we demonstrated previously in human osteosarcoma cells, expression of the RNA-DNA helicase Senataxin (SETX) in these cells reduced levels of single-strand breaks significantly (Figure 3C). Breaks were also eliminated by treatment of differentiated neuron-like cells with N-acetyl cysteine (NAC), an anti-oxidant (Figure 3C), or with ascorbic acid, another anti-oxidant (Figure S7).

Poly-ADP-ribosylation (PARylation) occurs at sites of DNA damage and we know that hyperPARylation is observed in the absence of ATM catalytic activity^11,32^. To quantify locations of PAR genome-wide we used ADP-ribose ChIP-seq^33^, a method of chromatin immunoprecipitation that employs a macrodomain specific for monomeric and polymeric forms of ADP-ribose^34^ in place of an antibody. Cells were differentiated for 7 days then crosslinked with formaldehyde and processed for ADP-ribose ChIP-seq. From this analysis we identified 2938 PAR peaks in the post-mitotic cells in the absence of any treatment, consistent with our observations of damage in untreated SH-SY5Y neuron-like cells^11^(Figure 3B,C). Examples of genome browser views of ADP-ribose ChIP-seq signal at several genomic locations are shown in Figure 4A. We also analyzed cells in which ATM activity was blocked with the ATM inhibitor AZD1390^35^ for the final two days of differentiation. Consistent with these observations, here we observed more PAR peaks (5028) in ATM inhibited cells compared to cells with normal ATM function.

**Figure 4.**
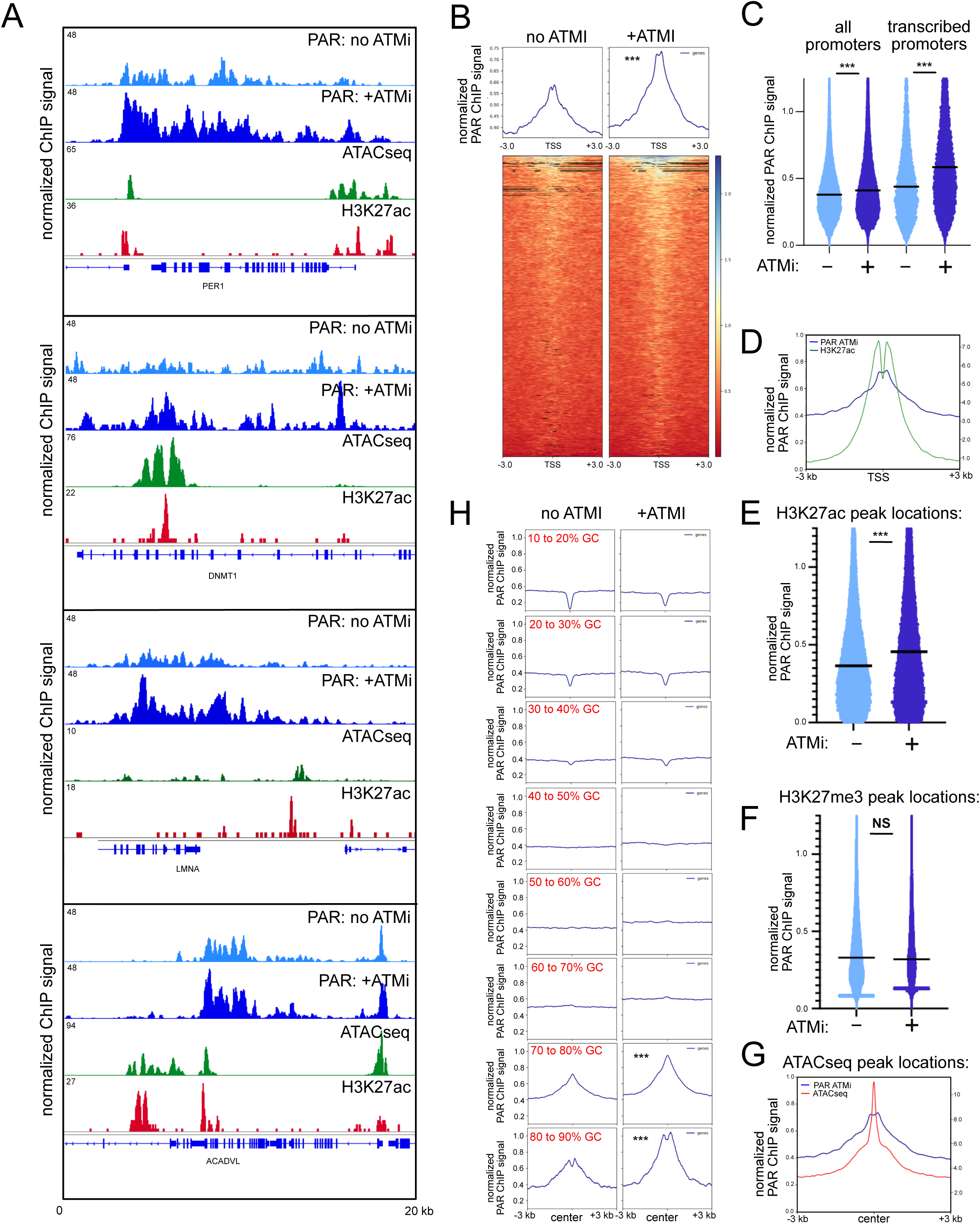
PAR ChIP shows increased signal with loss of ATM activity. (A) Examples of genome browser views of PAR ChIP in untreated SH-SY5Y differentiated neuron-like cells (“no ATMi”), cells with 2 day treatment with 1 μM AZD1390 (“+ATMi”), and ATACseq and H3K27ac data from previously published datasets^36,38^. (B) Recovery of PAR ChIP products in the presence or absence of ATMi; signal shown is ChIP signal from 2 replicates with input contribution removed and reads normalized by read depth for each sample. (C) Normalized PAR ChIP signal at all promoters (1000nt upstream of TSS to 500nt downstream) and promoters of genes that are transcribed, based on RNAseq data from SH-SY5Y cells. (D) Pattern of accumulated PAR ChIP signal in the presence of ATMi compared to the pattern of H3K27ac signal at TSS sites genome-wide. (E) Normalized PAR ChIP signal at H3K27ac peak locations (57,068). (F) Normalized PAR ChIP signal at H3K27me3 peak locations (176,439). (G) Pattern of accumulated PAR ChIP signal in the presence of ATMi compared to the pattern of ATACseq signal at ATACseq peak locations (“center”) genome-wide, including 3 kb upstream and downstream. (H) Normalized PAR ChIP signal is shown at subsets of ch1 locations with the indicated levels of GC content. 100nt sequence blocks (14,000 per bin for all bins except for 10 to 20 (13,181), 70 to 80 (13,677), 80 to 90 (2,523)) were used for each analysis. Locations of these blocks are shown (“center”) as well as 3 kb windows upstream and downstream. *** p<0.0005 by 2-sample t-test assuming unequal variances comparing untreated to ATMi, NS nonsignificant.

Our previous work also suggested that transcription stress is an important upstream factor in inducing ssDNA breaks and PAR. Our analysis here shows that 22% of all PAR peaks identified in untreated cells overlap a transcription start site (TSS) and this overlap increases to 35% in cells with ATM inhibition. Profiles of PAR signal at all TSS sites show a clear peak at the TSS that increases with ATM inhibition (Figure 4B). Consistent with this observation, we see higher normalized PAR ChIP signal at promoter-proximal regions (defined here as the region 1000nt upstream of the TSS through 500nt downstream) with ATMi exposure (Figure 4C). We used RNAseq data from differentiated SH-SY5Y cells to determine which genes are transcribed under these conditions and used this to identify the subset of expressed genes. When PAR ChIP signal is evaluated for these promoter regions specifically (“transcribed promoters”), higher levels of signal are observed, especially with ATM inhibition (Figure 4C).

The strong association with TSS sites suggests that PAR ChIP signal is likely correlated with chromatin marks found at active transcription locations. To analyze this we utilized previously published datasets for H3K27ac, H3K27me3, and ATACseq in SH-SY5Y cells^36–38^. The transcription-associated peaks in PAR signal align exactly in the center of the peak-valley-peak signal pattern of H3K27ac at promoters and enhancers (Figure 4D), predicted to be the nucleosome-free region at these active promoter and enhancer sites^39^. The accumulated PAR signal in ATM-inhibited cells at these H3K27ac sites is significantly higher than in untreated cells (Figure 4E), similar to the relationship at promoters. In contrast, no increase in PAR ChIP signal is observed at sites with H3K27me3, a repressive transcription mark (Figure 4F).

Active transcription marks are often coincident with measurements of “open” chromatin, as measured by ATACseq^40^. Previously published ATACseq data from SH-SY5Y cells^36^ also shows a correlation with the locations of PAR ChIP marks and levels of PAR with ATMi exposure are significantly higher at ATACseq peak locations compared to untreated neuron-like cells (Figure 4G). Nevertheless, there are also PAR ChIP signals in both untreated and ATMi-treated cells that appear to be independent of either ATACseq accessibility or H3K27ac marks (examples in Figure 4A).

Lastly, we analyzed the levels of PAR ChIP signal at genomic sites varying in GC content (100nt bins sampled across chromosome 1). This analysis shows that overall PAR is strongly biased toward high GC (> 70%) regions of the genome and that levels of PAR are significantly higher in these regions in ATMi-treated cells (Figure 4H). In contrast, we did not observe a positive correlation between PAR ChIP signal and GC skew (Figure S8).

### R-loops are coincident with PAR in differentiated SH-SY5Y neuron-like cells

We previously found that R-loops are higher in human cells depleted for ATM or treated with ATM catalytic inhibitors compared to untreated human cells^11^. In addition, the single-strand DNA breaks that appear with ATM loss as well as hyperPARylation can be reduced to normal levels with overexpression of Senataxin, an RNA-DNA helicase, suggesting that R-loops play a causal role in generating DNA damage in the absence of ATM function. If this is also the case in human neurons then we expect that locations of R-loops should be strongly correlated with locations of PAR. To test this idea we expressed a tagged catalytic mutant of human RNaseH in the neuron-like cells and used this to identify RNA-DNA hybrids, a method previously used in other cell types called “R-ChIP”^41^.

We observed R-loop peaks in both untreated as well as ATM inhibitor treated cells, and the levels of R-loops measured by R-ChIP in the differentiated neuron-like cells increased in the absence of ATM function, similar to our observations with PAR ChIP signal (examples of genome browser views in Figure 5A). This increase was eliminated in neuron-like cells simultaneously treated with the antioxidant N-acetyl cysteine (NAC), consistent with the idea that high levels of oxidative stress that form in the absence of ATM catalytic function are key to triggering these events. Importantly, the locations of R-loop signal were often overlapping with that of PAR ChIP signal in both untreated and ATMi-treated neuron-like cells. Nearly twice as many R-ChIP peaks were called in ATMi-treated cells (8018 peaks) compared to R-ChIP with ATMi and NAC exposure (4614 peaks).

**Figure 5.**
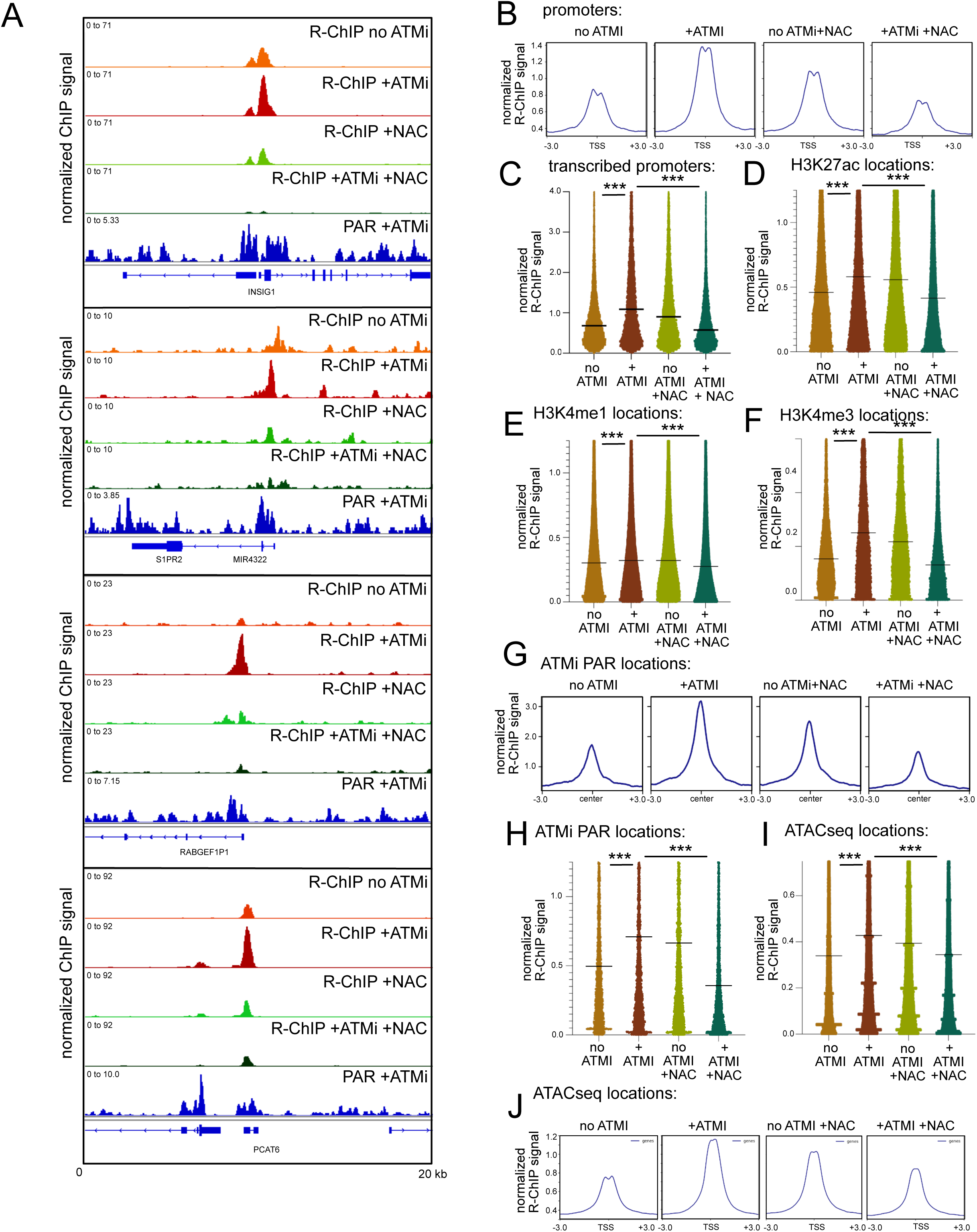
**R-loops increase with inhibition of ATM and are relieved by antioxidant treatment**. (A) Examples of genome browser views of R-ChIP in untreated SH-SY5Y differentiated neuron-like cells (“R-ChIP no ATMi”), cells with 2 day treatment with 1 μM AZD1390 (“R-ChIP +ATMi”), and cells with NAC treatment only (“R-ChIP +NAC), and cells with both ATMi and NAC (“R-ChIP +ATMi +NAC) as indicated in comparison to PAR ChIP signal in the presence of ATMi as in Fig. 4. (B) Pattern of accumulated R-ChIP signal at TSS sites genome-wide as indicated. (C) Normalized R-ChIP signal at transcribed promoters. (D,E,F) Normalized R-ChIP signal at H3K27ac, H3K4me1, and H3K4me3 ChIP locations, respectively^37^. (G) Pattern of accumulated R-ChIP signal at PAR ChIP sites in ATMi-treated cells genome-wide as indicated. (H) Normalized R-ChIP signal at PAR ChIP sites in ATMi-treated cells. (I) Normalized R-ChIP signal at ATACseq locations^36^. (J) Pattern of accumulated R-ChIP signal at ATACseq sites as indicated. *** p<0.0005 by 2-sample t-test assuming unequal variances.

As with locations of PAR, many R-ChIP peaks were also aligned with transcriptional start sites (61% of R-ChIP peaks measured with ATMi exposure overlap with at least one TSS). Quantification of R-loop peaks in all treatment conditions shows a 1.6-fold increase in total R-ChIP signal within 3 kb of TSS sites in ATMi-treated cells that is not observed in the presence of NAC (Figure 5B). This increase is clearly observed at transcribed promoters (Figure 5C). Similar patterns are also observed at H3K27ac, H3K4me1, and H3K4me3 locations that coincide with active promoters and enhancers (Figure 5D-F). Previous observations suggested that ssDNA breaks are primarily located at enhancers in human neurons^33^ but our analysis shows that a relatively small proportion (<2%) of R-ChIP peaks in neuron-like cells inhibited for ATM activity overlap with an enhancer, either ubiquitous or neuron-specific^42^.

Lastly, we examined the relationship between PAR ChIP locations and R-ChIP peaks and observed that a subset of R-ChIP peaks are localized coincident with PAR ChIP locations (Figure 5A,G). R-ChIP signal was approximately 2-fold higher in intensity at ATMi PAR ChIP locations in ATMi-treated cells compared with control cells or cells with both ATMi and NAC exposure (Figure 5G,H). As with PAR, the R-ChIP signal also closely aligned with ATACseq peaks (Figure 5I,J).

### PAR and R-loop patterns are aligned with transcription activity and GC content

The differences in PAR and R-ChIP patterns we observed suggested that transcription patterns may be altered in the cells with ATM inhibition. To examine this, we analyzed mRNA transcripts in SH-SY5Y differentiated cells in untreated, ATMi exposure, NAC, or ATMi plus NAC conditions using RNAseq. The results show minimal changes with ATM inhibition although NAC exposure induces significant alterations in transcription, both upregulated and downregulated genes (Figure S9).

Despite the lack of change in RNA transcripts with ATMi exposure in SH-SY5Y cells, we considered the possibility that the levels of PAR and R-loops observed at specific loci could be directly related to the level of transcription activity at these locations since our transcriptome analysis in A-T brain tissue showed a strong relationship between expression level in controls and the extent of change in A-T patients (Figure 2). To examine this question, we analyzed the mean PAR and R-ChIP signals in 100nt sliding windows across the genome, where each nucleotide position is represented by the mean ChIP signal between that position and 100nt upstream (Figure S10). Using this data, we determined the mean ChIP signal within the “promoter-proximal” region of every expressed gene (defined here as 1000nt upstream to 500nt downstream of a gene start for genes with corresponding RNASeq data) and compared these values to the corresponding transcript level for each gene (Figure 6A-C). This comparison shows a positive correlation between PAR and transcript abundance, particularly at high transcript levels (Figure 6A). A positive correlation between R-ChIP and transcript abundance is also observed that levels off above the lowest quartile of expression values (Figure 6B). Inhibition of ATM increased the overall PAR and R-ChIP signal but did not otherwise change the pattern.

**Figure 6.**
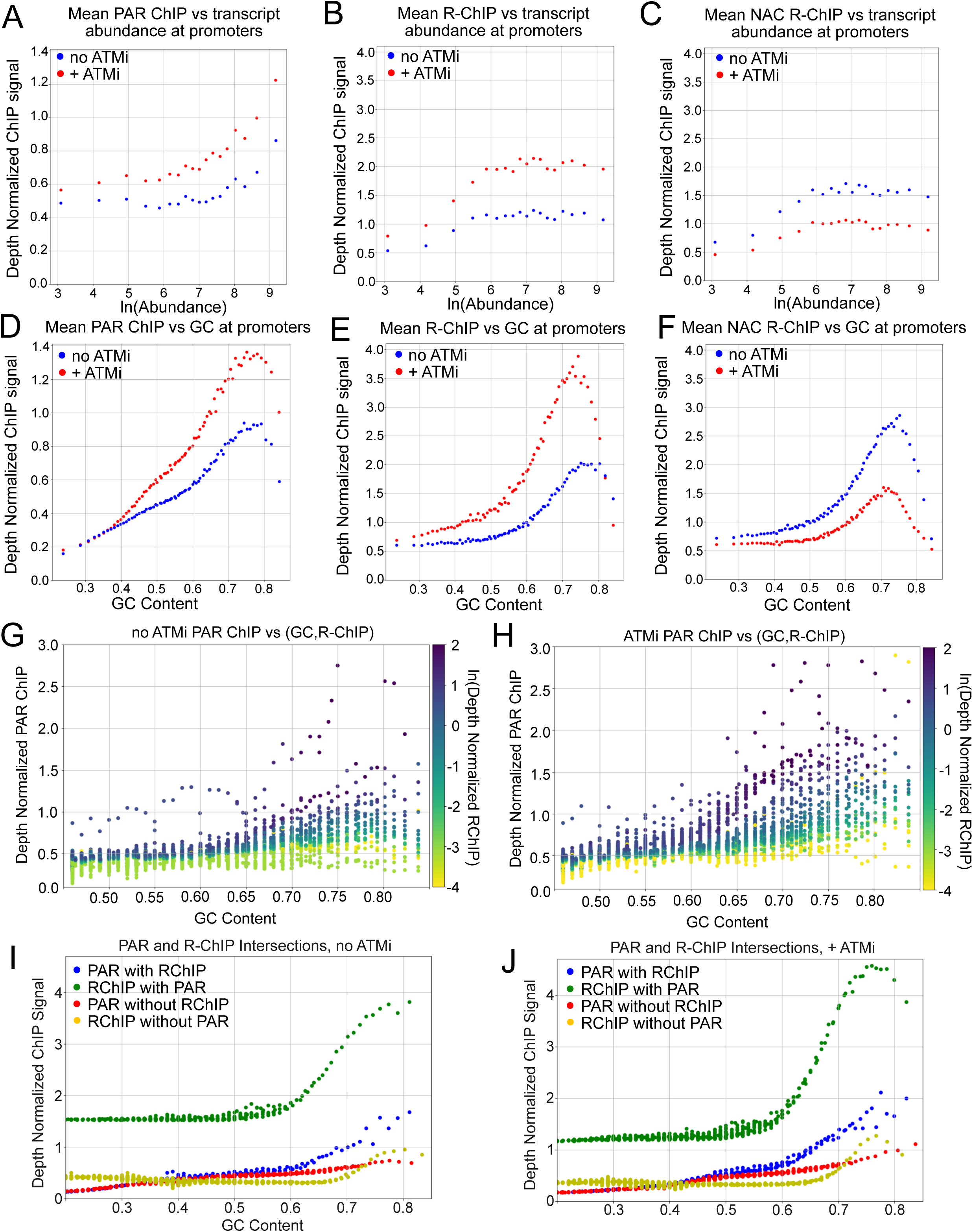
PAR and R-ChIP signals correlate with transcript abundance and GC content. (A) Mean TSS PAR ChIP plotted against ln(Abundance). TSS is defined in this figure as the window 1000nt upstream to 500nt downstream of a gene start. Sample space is comprised of locations where PAR is present within a TSS with corresponding transcript abundance info from RNASeq (“active TSS”)(See Fig. S10,S11). Sample space was sorted by transcript abundance value and divided into nonoverlapping bins of 1 million observations. The mean PAR and mean ln(abundance) value for each bin is plotted. Process was performed for no ATMi and + ATMi PAR samples. (B) As in (A) but for no ATMi and + ATMi R-ChIP. (C) As in (B) but for NAC treated R-ChIP. (D) Mean TSS PAR ChIP plotted against 100nt rolling mean GC content. Sample space comprised of locations where PAR is present within a TSS with corresponding transcript abundance info (“active TSS”). Sample was sorted by GC content and divided into nonoverlapping bins of 250K observations. The mean PAR and mean GC content for each bin is plotted. Process was performed for no ATMi and + ATMi PAR samples. (E) As in (D) but for no ATMi and + ATMi R-ChIP. (F) As in (E) but for NAC treated R-ChIP. (G) Mean PAR ChIP plotted against mean GC content and mean R-ChIP (no ATMi samples only). Sample space comprised of locations where PAR and R-ChIP signal are both present. Sample space was first sorted by GC content into nonoverlapping bins of 500K observations. Each 500K observation bin was then sorted by R-ChIP value before being divided into nonoverlapping bins of 25K observations. Each 25K bin was plotted based on mean PAR signal, mean GC content, and colored by ln(R-ChIP). (H) As in (G) but for + ATMi ChIP. (I) Mean ChIP signal per GC bins for various subsets of locations (no ATMi samples only). Subset of locations where PAR and R-ChIP coincide was sorted by GC content and divided into nonoverlapping bins of 250K observations. Subset of locations where PAR is present but R-ChIP is not and subset of locations where R-ChIP is present but PAR is not were each sorted by GC content and divided into nonoverlapping bins of 1 million observations. (J) As in (I) but for ATMi treated cells.

Consistent with the analysis of R-loop levels described above, NAC treatment reversed the relative R-ChIP signal patterns such that R-loops observed with ATMi and NAC treatment were lower than with NAC alone but showed an overall similar pattern relative to transcript abundance (Figure 6C).

Genome-wide GC content was calculated using the same 100nt sliding window scheme as the ChIP data (Figure S10, S11). The mean PAR and R-ChIP in promoter-proximal regions was plotted against GC content. Here we found that mean PAR signal is positively correlated with GC content with a maximum at approximately 75% GC (Figure 6D). Mean R-ChIP is also positively correlated with GC content but has a less dramatic increase until 50-60% GC and the highest values are also at approximately 75% GC, similar to PAR signal (Figure 6E). Analysis of PAR and R ChIP data with respect to GC skew showed higher levels overall with ATM inhibition, but much less dependence on the level of skew compared to GC content (Figure S8). As with the transcript abundance comparisons, ATMi treatment increased the overall signal of both PAR and R-ChIP but not the signal shape. With NAC exposure we observe the same overall GC dependence but the ATMi samples show dramatically lower levels of R-ChIP signal, below that of the untreated +NAC samples (Figure 6F).

When controlling for the GC influence on PAR, we observe that PAR signal in the control still correlates with R-ChIP signal (Figure 6G). If PAR and RChIP were coincidentally correlated because both correlate to GC content, this plot would not show a clear pattern of RChIP signal increasing as PAR increases for a given GC level. This is also observed in the ATMi sample but to a greater extent, with PAR bins separated by R-ChIP value extending higher, earlier when first binned by GC content (Figure 6H). To further characterize the relationship between PAR signal and R-ChIP, we compared the sets of all locations where PAR and R-ChIP coexist and exist separately. For both PAR and R-ChIP, signal was higher when the other was present and lower when separate (Figure 6I, 6J). These results show strong associations between PARylation and R-loop patterns that are not solely dependent on GC content.

## Discussion

In this work we identified and quantified R-loops and PAR-associated DNA damage sites in human post-mitotic neuronal-like cells with loss of ATM catalytic activity. The impetus for this was our previous study linking loss of ATM function to high ROS, transcriptional stress, and hyperPARylation^11^. Here, by mapping sites of PAR genome-wide, we have identified sequence patterns and correlations with histone marks that show that the PARylation events are strongly associated with gene promoters and high GC gene elements. These associations correlate well with patterns of transcriptional changes in A-T patients, as determined in this study by analysis of transcripts in cerebellum and cortex brain tissues.

### PARylation at transcription sites

We show in this and previous work^7,11^ that single-strand DNA breaks occur at a significantly higher level in ATM-inhibited neuron-like cells compared to untreated cells. This increase correlates well with the increases in PAR ChIP signal we have measured here. PARP1 and PARP2 are very rapidly activated at sites of DNA lesions to modify hundreds of proteins as well as undergoing automodification^43–45^. The PAR ChIP method was described by Nussenzweig and colleagues who showed that sites of PARylation in induced human induced neurons coincide with sites of DNA repair synthesis and that many of these are found at enhancer elements^33^. In parallel work, Gage et al also identified neuron-specific sites of repair-related fill-in synthesis and reported that these are associated with promoters, 5’ untranslated regions, and gene bodies^46^. Both studies found significant correlation between sites of DNA damage in neurons and regions of open chromatin mapped by ATAC-seq. Chromatin accessibility has also been found to be a key determinant of transcription disruption in a mouse model of cerebellar neurodegeneration with ATM and APTX loss^47^.

In our PAR ChIP data we see that 20% of called peaks in untreated neuron-like cells are located at promoter elements and 56% are associated with an annotated gene. With ATM inhibition the number and intensity of PAR ChIP peaks increases, consistent with the increase we observe in DNA damage by alkaline comet assays. 35% of PAR peaks called in ATMi-treated cells are located at promoter elements and 78% are overlapping with or within a gene. While we are largely equating PAR signal with DNA damage here, it is also possible that a subset of PAR ChIP signal we observe, particularly the low level basal signal, is associated with transcription independent of DNA damage since PARP1 is reported to be present at active genes and to regulate transcription in some cases^48,49^. Low levels of serine-targeted mono-ADP-ribose (MAR) were shown on chromatin throughout the cell cycle when the glycohydrolase ARH3 is eliminated^50^, perhaps an indication of “housekeeping”-associated, replication-independent MAR. This study by Ahel and colleagues also showed that excessive PAR production, as was observed in the absence of both ARH3 and PARG glycohydrolases, causes dysregulation of transcription.

### R-loops coincident with PARylation sites

Similar to our observations of PAR signal, we found that R-loops increased genome-wide with inhibition of ATM and that these sites were enriched in promoter regions as well as sites with active transcription marks. We used R-ChIP with catalytically inactive RNaseH here, which has the advantage that RNA-DNA hybrids are recognized within cells, before lysis and chromatin fragmentation^41^. Native recognition of R-loops in this way has been shown to best suited for quantification of R-loops formed by paused RNA polymerase in promoter-proximal regions^51,52^. The promoter-associated R-loops where we observe the largest changes with ATM inhibition are “class I” lesions, which have also been shown to be sites of R-loop accumulation in cells with dysfunctional splicing or BRD4^53,54^. Importantly, the increase in R-loop signals we observe with ATM inhibition is completely reversed with NAC antioxidant exposure, consistent with our previous observations using the S9.6 antibody and DRIP-qPCR^11^. These results show that the increase in promoter-associated R-loops caused by loss of ATM activity is dependent on the ROS generated under these conditions. Another example of ROS-induced R-loops comes from Lan and colleagues who use an inducible system to generate ROS at specific loci^55^. This work showed that R-loops are created in a ROS-dependent manner when induced at transcriptionally active sites and that these ultimately lead to transcription-coupled repair. Currently it is not known how ROS generates R-loops and whether stalling of RNA polymerase may contribute to this, or whether recruitment of repair factors to oxidative DNA damage may play a role.

Unexpectedly, we found that cells with normal ATM function appear to have higher levels of R-loops with NAC exposure. This increase also correlates with a large number of differentially expressed genes we observed in NAC-treated cells, similar to previous reports of NAC-treated cells in culture where this antioxidant is reported to be generally anti-proliferative^56^. Why NAC would increase levels of R-loops in normal cells is not clear, although it has been reported that the neutralization of ROS in otherwise untreated cells can reduce ROS to such low levels that it triggers “reductive stress”, an aberrant state involving increased mitochondrial oxidation and cytotoxicity^57^. Under these conditions, oxidative stress can actually be induced by reductive stress through feedback mechanisms^58^. Transcriptional upregulation in response to reductive stress could be the source of the increase in R-loops we observe.

### GC content

Our meta-analysis of PAR and R-ChIP signal, GC content, and transcription levels indicates that both PAR and R-loops are strongly correlated with gene expression levels as well as with DNA sequence. GC content and GC skew have been known to be associated with R-loop propensity for many years and several classes of promoter region have been defined in mammalian cells based on the levels of R-loops and sequence context^51,59^. Here we find that sites with high GC content overall have PAR and R-loops, particularly with ATM inhibition, and that the sites that exhibit the highest signals show both marks. Interestingly, there appears to be a threshold of GC content that promotes a transition to high PAR/R-loop propensity. Genomic sites with GC content above this level, approximately 60%, delineates groups of loci that are associated with both marks.

### Gene expression levels

In SH-SY5Y-derived neuron-like cells with ATM inhibition there are relatively few changes in gene expression compared to untreated cells despite the accumulation of PAR and R-loops that we demonstrate here. In contrast, we found dramatic changes in transcripts in brain tissue from A-T patients compared to controls. The pattern of downregulated genes in the cerebellum closely matches the GC content trends seen in the neuron-like cells in culture, however, and the most affected genes had the highest level of expression in control individuals, similar to the expression patterns we observed with PAR and R-ChIP signals in cells. We hypothesize that the loss of transcripts from highly-expressed genes in A-T cerebellum likely requires a significant duration of ATM loss, especially considering the progressive nature of cerebellar atrophy in A-T and the ages of the patients analyzed here (up to 31 years).

A-T patients show a loss of cerebellum volume over time which is visible in CT and MRI images^30^. These changes are often not apparent early in the course of the disease when functional changes are starting; however, MRI studies do show a progressive cerebellar atrophy in patients over time^1^. Loss of Purkinje cells from the cerebellum is known to be characteristic of A-T patients, and the importance of this cell type for the function of the cerebellum suggests that the disappearance of Purkinje cells likely is related to loss of cerebellar function. Here we considered the possibility that loss of Purkinje cells may be responsible for the apparent loss of highly-expressed transcripts simply because the cells are absent in the patients. We determined the density of Purkinje neurons along the Purkinje cell layer of A-T patients and control individuals and found that, as a group, there is a reduction in Purkinje cell density in patients of approximately 40%. Given the magnitude of this change, it is possible that transcripts reduced by 30 to 50% may result from the partial loss of this cell type. A subset of the downregulated transcripts fall into this category, including ITPR1 and CA8, genes with expression loss at the protein level that we observed by mass spectrometry previously^11^. Several hundred gene transcripts are reduced more than 4-fold in A-T cerebellum, however, so these are not likely to be lower simply due to loss of this cell type. Another piece of evidence in this regard is the fact that over 800 transcripts are downregulated in the frontal cortex tissue which does not lose significant tissue mass or cell types in A-T^30,60^. The characteristics of downregulated genes in the cortex match the GC content and expression patterns of genes affected in the cerebellum, which argues that the patterns are due to transcriptional changes resulting from loss of ATM rather than the loss of a specific cell type in the tissue. It is not known why ATM loss affects the cerebellum more than other parts of the brain although it is clear that there is a cerebellum specificity to many DNA repair-related syndromes that involve single-strand DNA breaks^12^. In addition, the cerebellum shows some of the highest levels of ROS production compared to other brain regions^61^.

In addition to Purkinje cell-specific transcripts, we also observed a large number of other changes in gene expression that were significant even when considering the magnitude of Purkinje cell loss in the tissue. While A-T patients are known to lose Purkinje neurons, it is likely that abnormalities in other cells in the cerebellum are also critical. It is worth noting that concomitant loss of *ATM* and *POLB* in the mouse leads to cerebellar ataxia but not to loss of Purkinje cells^62^, thus it is possible that other events are important drivers of neurodegeneration outside of this cell type. For instance, several groups have demonstrated effects of ATM loss on microglia that suggest that these immune cells play very important roles in regulating neuroinflammation and neuronal function^63–65^.

Overall, the results shown here illuminate the genome-wide patterns of DNA damage and R-loops in neuron-like cells lacking ATM function and support the hypothesis that these molecular events may underlie ROS-driven changes in gene expression. The transcriptional changes we observed in A-T patient cerebellum tissue are very likely related to the pathology of the disease considering that many genes implicated in cerebellar ataxia are significantly reduced in patient samples. Further work is still needed to determine the source of R-loop-associated single-strand DNA breaks and to generate a biological model system with human cells to study the progression of damage and how it affects transcription patterns over time.

## Acknowledgments

We thank members of the Paull laboratory for helpful discussion and Vishwanath Iyer for useful suggestions. We are indebted to the families of the A-T patients and control individuals who contributed the autopsy material for research use, as well as the Neurobiobank for providing us with the frozen and fixed tissue samples. We acknowledge grant support from the A-T Society, NIH R01NS126747, and Cancer Prevention and Research Institute grant RP200254. Sequencing was performed by the UT Genomic Sequencing and Analysis Facility (RRID:SCR_021756).

## Author Contributions

Conceptualization, J-H.L., T.T.P, P.R.W.; Methodology, J-H.L., X.W., O.M.C., P.R.W., N.A.E.; Investigation, J-H.L., X.W., O.M.C., P.R.W., N.A.E., T.T.P.; Writing original draft, T.T.P., P.R.W.; writing review and editing, J-H.L., X.W., O.M.C., P.R.W., N.A.E., T.T.P.; Funding acquisition, T.T.P; Supervision, T.T.P., X.W.

## Materials and Methods

### Cell culture

Human SH-SY5Y neuroblastoma cells obtained from ATCC (CRL-2266) were grown in DMEM (Invitrogen) supplemented with 10% fetal bovine serum (FBS, Invitrogen) and 100 units/ml penicillin-streptomycin (15140-122, Life Technology), adherent cells only. The cells were induced to differentiate using a 3 day treatment with 1% FBS media and 10 μM retinoic acid (Sigma R2625) followed by 4 days with zero FBS, 10 μM retinoic acid, and 50 ng/ml BDNF (R&D Systems 248-BDB-250). Human RNaseH was expressed using a lentivirus derived from pTP5135, containing a tet-on allele of RNaseH D210N in a pLenti Hygro H2B mRuby backbone. pLentiPGK Hygro DEST H2B-mRuby2 was a gift from Markus Covert (Addgene plasmid # 90236; http://n2t.net/addgene:90236; RRID:Addgene_90236)^66^. Lentivirus was prepared in HEK-293T cells as previously described ^7^; SH-SY5Y cells were transduced with lentivirus and selected in hygromycin (200μg/ml). For experiments with ATM inhibitor, AZD1390 (Selleckchem S8680) was added for the final two days of differentiation at a concentration of 1 μM). For experiments with NAC, 1 mM was added for the final 2 days of differentiation.

### Comet Assays

Comet assays were performed using the Cell Biolabs OxiSelect comet assay kit according to manufacturer instructions with the following modifications: The gel was run using a Horizon 11-14 agarose gel apparatus using 35V for 15 to 30 min. for alkaline comet assays and 50V for 15 min. for neutral comet assays. Images were obtained using a Nikon TS2-FL microscope and analyzed with OpenComet^67^.

### ROS measurements

CM-H2DCFDA (Thermo Fisher C6827) ROS assays were performed with differentiated SH-SY5Y cells using 1 μM CM-H2DCFDA for 15 min and analyzed by FACS.

### PAR ChIP

The protocol was adapted from ^33^. Briefly, SH-SY5Y cells were differentiated as described above and two 15 cm dishes were used for each ChIP replicate (approximately 20M cells). Cells were removed from dishes using trypsin and then resuspended in media in a volume of 36 ml.

Formaldehyde (37%, Sigma F1635) was added to 1% final concentration and incubated at 37°C for 7 min. with occasional inversion of the tube. 2.5M glycine was added to a final concentration of 0.125M. The tube was inverted several times and cells recovered by centrifugation at 550xg for 5 min. at 4°C. The pellet was washed twice with 10 ml cold PBS using 550xg centrifugation for 5 min. each time. The pellets were snap frozen using liquid nitrogen and stored at –80°C for later use. Each cell pellet was resuspended in 1mL cold PAR ChIP RIPA buffer (10 mM Tris-HCl, pH7.6, 1 mM EDTA, 0.1% SDS, 0.1% sodium deoxycholate, 1% Triton X-100) containing protease inhibitor (Fisher, A32955). Samples were sonicated using a Diagenode Bioruptor UCD-200 instrument on high power with 15 sec on/ 15 sec off cycles for 45 min. at 4°C. Under these conditions, fragmented chromatin was 200 to 500 bp. Samples were centrifuged at 20,000xg for 10 minutes at 4°C. The supernatant was transferred to another tube; 50 μl were reserved as input DNA. 40 μl of Protein A/G magnetic beads was incubated with 5 μg anti-pan-ADP-ribose binding reagent (Millipore Sigma, MABE1016) per IP sample in 100 μl PBS for 30 min. then washed with 200μl PBS twice. The beads were then added to the supernatant and incubated at 4°C overnight. The following day, beads were washes for 10 min. at 4°C with the following sequence of buffers: 2x with 1 mL of RIPA buffer, 2x with 1 mL of RIPA buffer + 0.3M NaCl, 1x with 1 mL of TE pH 8 + 0.2% Triton X-100, 1x with 1 mL of TE pH 8. The beads were resuspended in 100μl TE; 3μl 10% SDS and 5μl 20 mg/ml proteinase K were added and the samples incubated at 65°C for 4 hrs while rocking. The tubes were vortexed every 30 to 50 min. The DNA was removed from beads using a magnetic stand; 100μl TE + 0.5M NaCl was added to the beads and then removed and combined with the first sample. 200 μl phenol/chloroform/isoamyl alcohol was added and samples were added to Phase Lock gel columns (VWR, 10847-802) and processed according to manufacturer instructions. Samples were then ethanol precipitated by adding 2μl glycogen (20 mg/ml), 20μl 3M NaOAc pH 5.2, and 500 μl cold ethanol (100%). After incubation on dry ice for 15 min., samples were centrifuged at 20,000xg for 15 min. at 4°C. Pellets were washed with 70% ethanol twice; the supernatant was removed and pellets dried at room temp. Pellets were resuspended in 22 μl 10 mM Tris-HCl pH 8.0, vortexed, and stored at –20°C.

### R-ChIP

The protocol was adapted from ^41^. Briefly, SH-SY5Y cells were differentiated as described above and two 15 cm dishes were used for each ChIP replicate (approximately 20M cells). Cells were crosslinked in the dishes with the addition of formaldehyde to 1% final concentration and incubated for 7 min. while shaking (120 RPM). 2.5M glycine was added to a final concentration of 0.125M. Cells were removed by scraping and transferred to conical tubes with an additional 5 ml PBS to wash the dishes. Cells were centrifuged at 2187xg for 15 min. at 4°C, then washed with cold PBS and pelleted again. The pellets were snap frozen using liquid nitrogen and stored at –80°C for later use. Each cell pellet was resuspended in 2.3 mL cold R-ChIP RIPA buffer (50 mM Tris-HCl, pH 8.0, 0.15M NaCl, 2 mM EDTA, 0.1% SDS, 0.5% sodium deoxycholate, 0.1% Igepal CA-630 (NP40) containing protease inhibitor (Fisher, A32955) and incubated on ice for 10 min.

Samples were sonicated using a Diagenode Bioruptor UCD-200 instrument on high power with 15 sec on/ 15 sec off cycles for 45 min. at 4°C. Samples were centrifuged at 3,889xg for 10 minutes at 4°C. The supernatant was transferred to another tube; 50 μl were reserved as input DNA. 25 μl of Protein A/G magnetic beads per IP sample were washed with 1 ml wash buffer (20 mM Tris-HCl pH 8.0, 0.15M NaCl, 2 mM EDTA, 1% Triton X-100) using a magnetic separator; this step was repeated for a total of 3 washes. Beads were resuspended in bead blocking buffer (0.2 μg/ml BSA, 0.2 mg/ml glycogen in PBS) and incubated for 1 hr at RT. Beads were then washed 3 times with antibody binding buffer (5 μg/ml BSA in PBS), resuspended in 250 μl antibody binding buffer and 2.5 μg FLAG M2 antibody (Sigma) per 25 μl of Protein A/G magnetic beads and incubated overnight at 4°C with rotation. After removal of unbound antibody, beads were resuspended in 1 ml sonicated lysate and incubated overnight at 4°C with rotation. The following day, beads were washed 3 times for 3 min. at room temperature with 1 ml wash buffer I (20 mM Tris-HCl pH 8.0, 150 mM NaCl, 2 mM EDTA, 1% Triton X-100, 0.1% SDS) containing protease inhibitor, then 3 times with wash buffer II (wash buffer I containing 0.5M NaCl), then once with wash buffer III (20 mM Tris-HCl pH 8.0, 1 mM EDTA, 1% NP40, 0.25M LiCl), then once with TE. Samples were moved to a new tube and eluted with 100 μl elution buffer (0.1 M sodium bicarbonate, 1% SDS) at 30 °C for 30 min. Beads were removed, 1 μl RNase A (20 mg/ml) was added to each sample and supernatants were decrosslinked at 65°C for 22 hrs. 1 μl

Proteinase K (20 mg/ml) was added to each sample and incubated for an additional 2 hrs at 65°C. Samples were purified using the nucleotide removal kit (Qiagen) according to manufacturer instructions.

### ChIP library preparation and sequencing

For PAR and R-ChIP, the eluted DNA (2 biological replicates per condition) as well as input samples were used to make sequencing libraries using the NEBNext Ultra II DNA Library Prep Kit for Illumina (NEB) with NEBNext Multiplex dual index primers using 12 amplification cycles and 2 additional AMPure XP clean-up steps at 0.8X. Libraries were sequenced by the UT Genomic Sequencing and Analysis Facility (RRID:SCR_021713) using the Illumina NovaSeq SP platform with PE150 runs.

### RNA extraction from patient tissues

Fresh-frozen A-T patient and control tissues were obtained from the NIH Neurobiobank (Table S1). 50mg patient tissue RNA was extracted in an RNase free environment using RNA Extraction kit for Bioruptor Plus (Diagenode Cat# C20000010). The additional protocol listed for DNaseI treatment was followed for 10ug of extracted RNA per sample. DNaseI treated RNA was cleaned using RNeasy Mini Kit (Qiagen Cat# 74104). RNAseq library prep was performed using NEBNext Ultra II RNA Library Prep for Illumina (NEB Cat# E7770S). The NEB protocol was followed using the Poly(A) mRNA Magnetic Isolation Module and SPRIselect beads (Beckman Coulter Cat #B23317). Library adapter was diluted 5-fold. The additional SPRIselect cleanup step referred to at 1.11.3 in the NEB protocol was followed.

### Purkinje cell density estimation

Purkinje cell counts were done in triplicate for each A-T patient and control using previously generated chromogenic immunohistochemistry image files ^11^. For each image, Purkinje cells were enumerated and normalized by the total length in μm of the Purkinje layer as measured in Fiji.

### RNA Extraction from SH-SY5Y neuroblastoma cells

Total RNA extraction was performed using NEB Monarch Total RNA Miniprep Kit (New England Biolabs Cat #T2010S). RNAseq library prep was performed using NEBNext Ultra II RNA Library Prep for Illumina (NEB Cat# E7770S). The NEB protocol was followed using the Poly(A) mRNA Magnetic Isolation Module and SPRIselect beads (Beckman Coulter Cat #B23317). Library adapter was diluted 5-fold. The additional SPRIselect cleanup step referred to at 1.11.3 in the NEB protocol was followed.

### RNAseq computation

Sequencing reads were cleaned using FASTP in paired end mode, transcript abundance levels were determined using Salmon pseudoaligner and GRCh38 (hg38), and differentially abundant transcripts were quantified using Swish in the Gene mode. Statistically significant differences between density plots were determined using empirical cumulative distribution function tests via DTS R package with 5000 bootstraps. GSEA information was computed via the Fast Gene Set Enrichment Analysis (FGSEA) R package.

### ChIP computation

Sequencing reads were cleaned using FASTP in paired end mode and mapped to the genome using STAR aligner and GRCh38 (hg38). Aligned reads were converted to bedgraph format via MACS2 ^68^. Output bedgraph files were processed again using MACS2’s bdgcmp command with the ppois flag to approximately remove the ChIP input influence from the ChIP bedgraph files. Finally, bedgraph output was read depth normalized to make files equivalent in total signal.

For analysis shown in Figures 4 and 5, bedgraph files were converted to bigwig format and analyzed using deepTools computematrix and plotheatmap or with multibigwigsummary ^69^. Bed files for comparisons at specific sites were taken from refTSS^70^, data from this study, or from previously published GEO datasets SRR3363259, SRR3363257, SRR3363256, SRR3363255, SRR3363258, SRR5819663, SRR5819664 ^36–38^. Processing of external data was done as described for the other raw data; bed files used for comparisons were peaks called using MACS2 with input files as controls.

For analysis shown in Figure 6, Final bedgraph files were separated into separate bedgraph files for each chromosome. Bedgraph files were converted into “extended” Numpy arrays where each array element represents a nucleotide on the genome and the element value is the bedgraph value at that location. Pandas *rolling(window=100)* function was paired with *mean()* to calculate the 100nt rolling mean ChIP signal across the genome (see Figure S10). Rolling mean GC Content was computed similarly. GRCh38 genome sequence was converted to integer data type (0 for A/T and 1 for G/C) before the 100nt rolling mean was computed as above. Processed data using these arrays for figures was all computed using Numpy and Pandas. Plots were made using Matplotlib.

### Visium spatial transcriptomics

Fresh-frozen A-T and control tissues (A-T patient 1 and Control 2, Table S1) were sectioned, mounted on Visium slides, and sequenced using a NovaSeq 500 platform with PE150.

### Visium Preprocessing

Spaceranger software was used to align dot transcription information onto tissue slice stains. Aggregation was performed to merge the AT and Control samples into one file.

Actual commands:

spaceranger count ––id=836_AT \

––transcriptome=/stor/work/Paull/refdata-gex-GRCh38-2020-A \

––fastqs=/stor/work/Paull/paull_bcl/outs/fastq_path/ \

––sample=836AT \

––image=/stor/work/Paull/tifs/836AT.tif \

––unknown-slide \

––localcores=8 \

––localmem=64 \

––reorient-images spaceranger count ––id=5829_C \

––transcriptome=/stor/work/Paull/refdata-gex-GRCh38-2020-A \

––fastqs=/stor/work/Paull/paull_bcl/outs/fastq_path/ \

––sample=5829C \

––image=/stor/work/Paull/tifs/5829C.tif \

––unknown-slide \

––localcores=8 \

––localmem=64 \

––reorient-images spaceranger aggr ––id=836AT_5829C \

––csv=aggregate.csv \

––normalize=mapped

Once quantified, the Cloupe files were analyzed using 10X Genomics Loupe Browser 6. Cell layers were manually assigned using the stain layers and candidate gene markers as reference. tSNE plots were processed using Loupe Browser 6 as well. Once linking each dot to a cell layer, transcript counts for each gene in each dot was exported using instructions from this 10X tutorial: https://kb.10xgenomics.com/hc/en-us/articles/360023793031-How-can-I-convert-the-feature-barcode-matrix-from-Cell-Ranger-3-to-a-CSV-file-?source=search. We fed dot transcript data into DESeq2 where each dot was considered a replicate for each condition (AT or C). Purkinje cell markers were downloaded from an annotated, developing cerebellum scRNAseq dataset provided by UCSC linked here: https://cells.ucsc.edu/?ds=cbl-dev.

### GC Skew

GC Skew was calculated as (#G-#C)/(#G+#C).

### Poisson Comparison of ChIP data

Statsmodels (v 0.14.0) Poisson ratio test of independence was used to compare mean ChIP values between samples. Sample sizes were 1,000 to save on computational resources.

Poisson parameters for each distribution were the mean values plotted.

Actual command:

test_poisson_2indep(1_000, Mean1, 1_000, Mean2, method=’etest-score’,compare=’ratio’)

### Data availability

All sequencing data is available via the NIH GEO database (GSE233479). All code generated for this project is available at https://github.com/prwoolley/ATM_par_rchip_paper.

