## Supplemental Figures for "Regulation of transcription patterns, poly-ADP-ribose, and RNA-DNA hybrids by the ATM protein kinase"

Figure S1: GSEA analysis of transcripts dysregulated in A-T cerebellum tissue

Figure S2: analysis of Purkinje cell numbers in A-T cerebellum tissue

Figure S3: Analysis of cerebellum transcript data for relationships with GC skew

Figure S4: Spatial transcriptomic analysis of a control cerebellum tissue and an A-T patient tissue.

Figure S5: ROS measurements in differentiated SH-SY5Y cells.

Figure S6: Neutral comet assays with ATM inhibition

Figure S7: Alkaline comet assays with ATM inhibition and ascorbic acid antioxidant

Figure S8: Analysis of PAR ChIP and R ChIP data for relationships with GC skew

Figure S9: RNAseq analysis of SH-SY5Y neurons with ATM inhibition

Figure S10: Rolling mean ChIP analysis strategy

Figure S11: ChIP sample space histograms

### Supplemental Tables

Table S1: A-T patient and control tissues used in Purkinje cell counting and RNAseq analysis

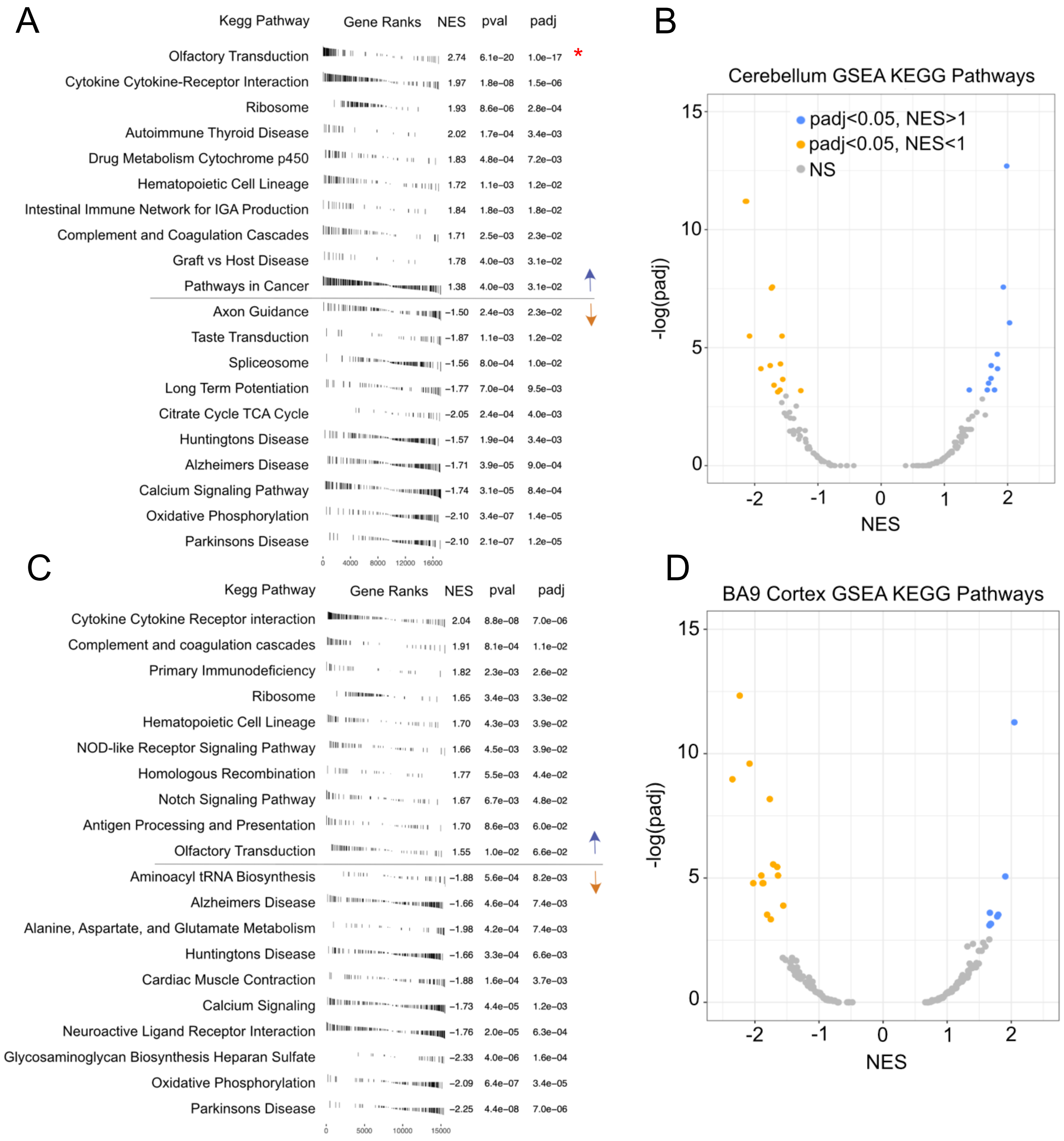


**Figure S1. GSEA analysis of dysregulated KEGG pathways in A-T brain tissue.**
(A) KEGG pathways showing significant differences between A-T and control cerebellum tissues. Genes were ranked based on log2FC before GSEA analysis(1). The 10 pathways with the lowest adjusted p-value for the positive normalized enrichment score set (NES>0) are displayed on the top (blue arrow). The 10 pathways with the lowest adjusted p-value for the negative normalized enrichment score set (NES<0) are displayed on the bottom (orange arrow). (B) Volcano plot of GSEA analysis showing all KEGG pathways identified in cerebellum data, with significant upregulated (blue) and downregulated (orange) pathways indicated. Adjusted P-values (Y axis) compared to NES scores are shown, with the exception of Olfactory transduction not shown here. (C) As in (A) but for the BA9 cortex. (D) As in (B) but for the BA9 cortex.

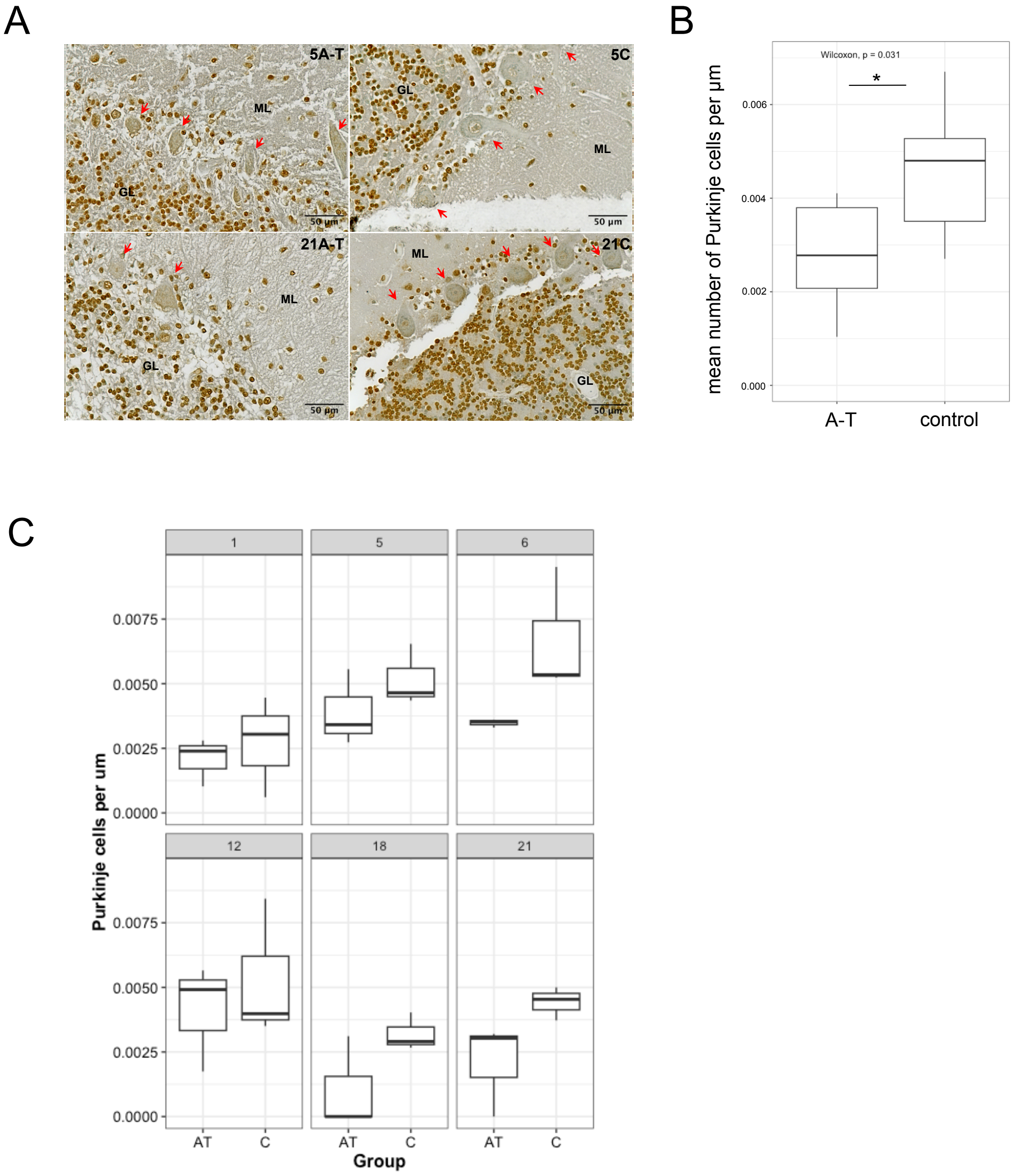


**Figure S2. Reduction of Purkinje cell numbers in A-T patients is variable.** (A) Examples of cerebellum tissue images for control and A-T individuals with Purkinje cells marked in red. (B) Purkinje cell counts from 6 A-T patients and 6 controls using previously generated chromogenic immunohistochemistry image files (2). Mean Purkinje cell counts were significantly less (~40%) in the A-T patient group as compared to the control group (p=0.031 by Wilcoxon test for paired samples). (C) Purkinje cell counts for each A-T patient and their age-matched control. Differences for each pair individually were not statistically significant by Wilcoxon test.

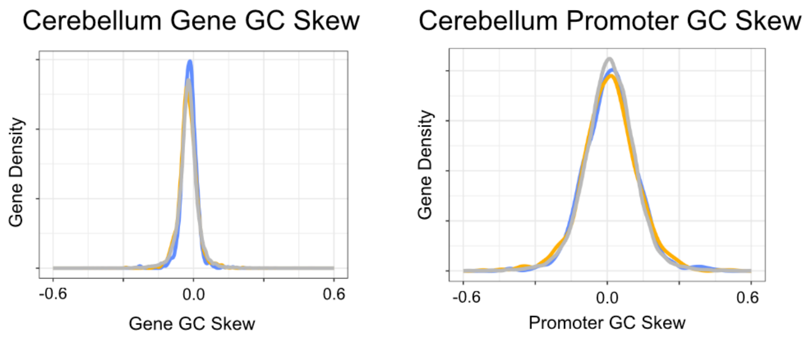


**Figure S3. Analysis of cerebellum transcript data for relationships with GC skew.** Transcripts observed to be lower (gold), higher (blue) or unchanged (grey) in A-T cerebellum compared to wild-type cerebellum samples are shown with the level of GC skew compared to transcript abundance. GC skew is defined here as $(G - C)/(G + C)$ and is calculated for the entire gene (left) or for the promoter region only (right, 1000nt upstream of TSS plus 500 nt downstream of TSS).

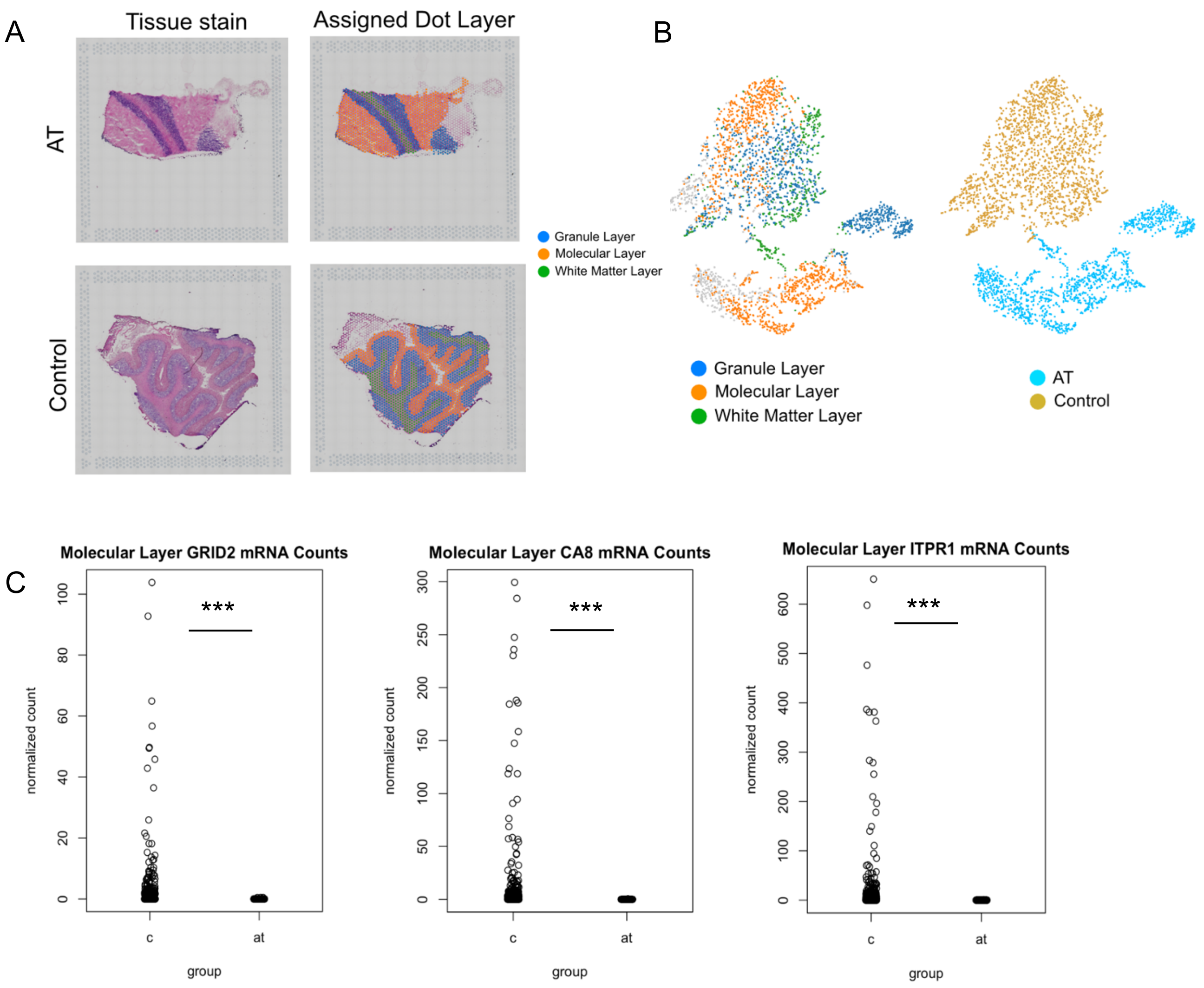


**Figure S4. Spatial transcriptomic analysis of a control cerebellum tissue and an A-T patient tissue.** (A) Layer assignment for 10X Visium dots. Left panels show hematoxylin and eosin stained tissue slices. Right panels are of the assigned cell layer for each dot (molecular, granule, or white matter). Dot assignment was informed by stained tissue images which show distinct separation of three layers as well as dot clusters determined by gene markers using Loupe Browser 6 (software provided by 10X Genomics for Visium data). Each dot is 55 μm, placed in a grid with approximately 100 μm between dots. Each dot contains oligonucleotides with poly(dT) for capture of poly(A) mRNA, a unique spatial barcode, a unique molecular identifier, and a sequencing primer recognition sequence. (B) tSNE for each dot across the A-T and control tissue slices. The tSNE plot demonstrates generally good cell layer separation from transcript data in the Control tissue, validating our manual assignment of cell layers for each dot. Additionally, there was complete separation of the A-T tissue slice from the control tissue slice, demonstrating little transcription similarity between cells of the same layer between A-T and Control, for all cell layers. (C) Examples of DESeq2 normalized gene counts for Purkinje marker genes in molecular layer dots between A-T and Control tissue slices. All dots from the molecular layer in each tissue slice were fed into DESeq2, where each dot from a respective tissue slice was treated as a replicate and all dots from the AT group was compared against the Control group. Note that not all molecular layer dots contain Purkinje cells, hence many Control tissue dots having low transcript counts for these genes. *** indicates $p<0.001$ generated by DESeq2 using the Wald test for hypothesis testing when comparing two groups.

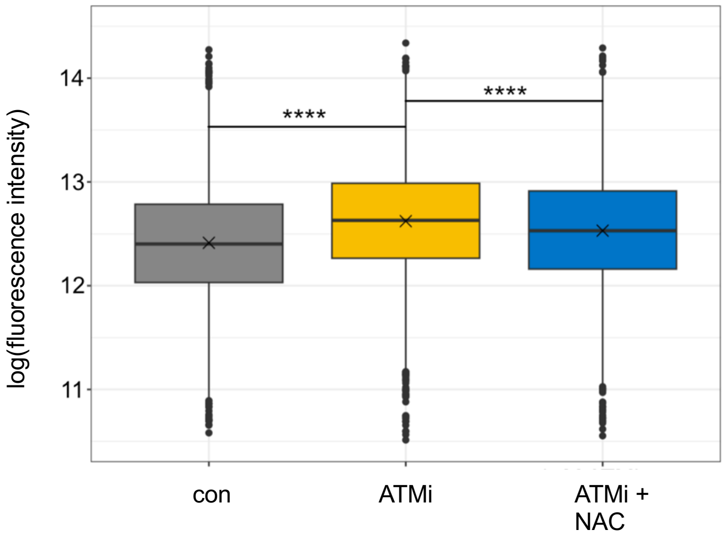


**Figure S5. ATM inhibition results in higher ROS.**
Total ROS levels were measured by FACS using CM-H2DCFDA in differentiated SH-SY5Y neuronal-like cells that were untreated (con) or treated with 1 μM AZD1390 (ATMi) or ATMi with 1 mM NAC (ATMi + NAC) during the final two days of differentiation. Line indicates median. **** $p<0.0001$ by 2-sample t-test assuming unequal variances, NS= non-significant.

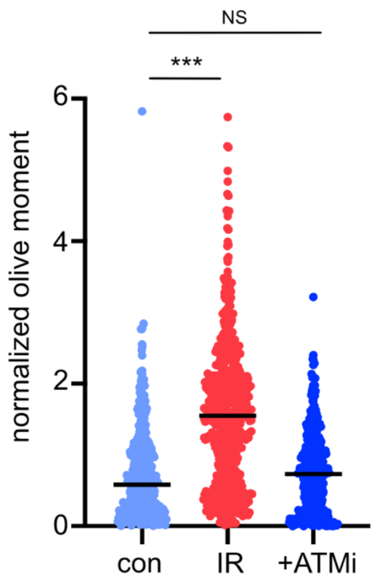


**Figure S6. ATM inhibition does not induce DNA double-strand breaks in differentiated neuron-like cells.** Neutral comet assays were performed in differentiated SH-SY5Y cells with >350 cells per group. Cells were either untreated (con), treated with 20Gy ionizing radiation (IR), or treated with 1 μM AZD1390 (ATMi). Line indicates median. *** $p<0.0005$ by 2-sample t-test assuming unequal variances, NS= non-sigificant.

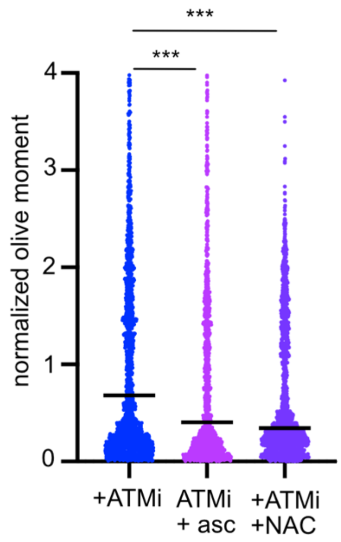


**Figure S7. Ascorbic acid reduces single-strand DNA breaks in ATM-inhibited neuron-like cells.** Alkaline comet assays were performed in differentiated SH-SY5Y cells with >350 cells per group. Cells were either treated with 1 µM AZD1390 only (ATMi), ATMi with 100 µM ascorbic acid (ATMi + asc), or ATMi with 1 mM NAC (ATMi + NAC). Line indicates median. *** $p<0.0005$ by 2-sample t-test assuming unequal variances.

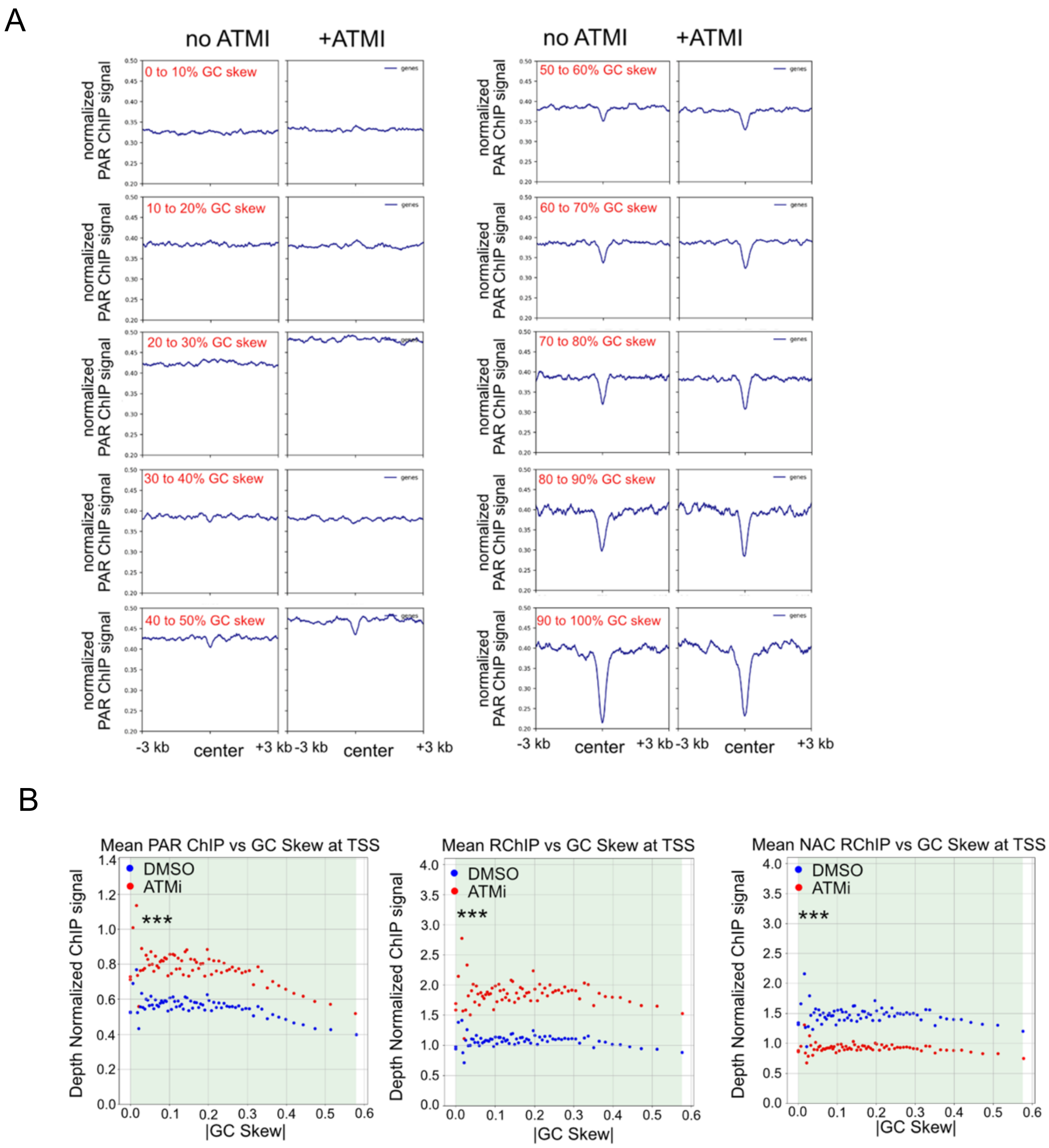


**Figure S8. Analysis of PAR ChIP and R ChIP data for relationships with GC skew.** (A) PAR ChIP signal from cells with no treatment or cells with ATM inhibitor treatment was analyzed with respect to GC skew. GC skew is defined here as *(G − C)/(G + C*) and is divided into 10 bins of genomic regions with the indicated range of GC skew. 14,000 randomly chosen 100nt genomic locations from ch1 are used for this analysis per bin, except for 70 to 80% (6650 regions), 80 to 90% (2864 regions), and 90 to 100% (3547 regions). (B) PAR ChIP and R-ChIP signal from cells with no treatment or cells with ATM inhibitor treatment and with or without NAC treatment is analyzed with respect to GC skew, defined as in (A). Green shaded regions indicate locations where the no ATMi PAR signal is significantly different than the + ATMi PAR signal ($p<0.001$). Significance was determined by a Poisson ratio test of independence using the statsmodels “test_poisson_2indep()” function. The Poisson parameter was the mean PAR signal for each bin.

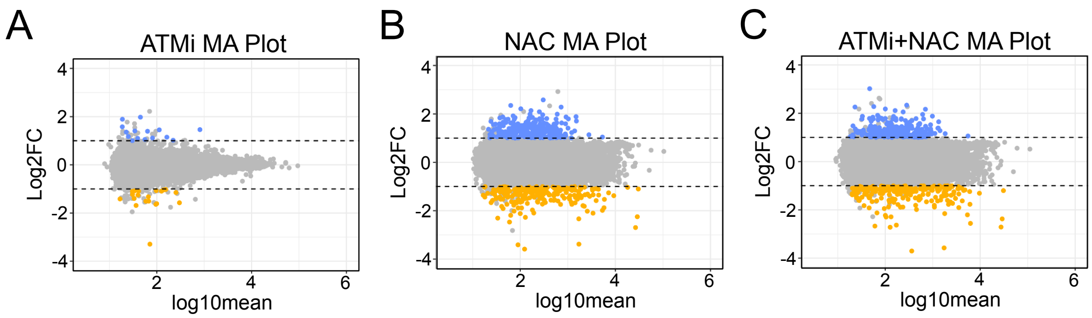


**Figure S9. RNAseq analysis of SH-SY5Y neurons with ATM inhibition.** (A) MA plot comparing ATMi-treated SH-SY5Y differentiated neurons to untreated cells. Dashed lines correspond to |Log2FC|>1. Orange corresponds to Log2FC<-1 with qvalue<0.01 (downregulated gene set) and blue corresponds to Log2FC>1 with qvalue<0.01 (upregulated gene set). (B) MA plot comparing NAC-treated SH-SY5Y differentiated neurons to untreated cells. (C) MA plot comparing ATMi + NAC-treated SH-SY5Y differentiated neurons to untreated cells. ATMi (AZD1390) was added for the final two days of differentiation at a concentration of 1 μM). For experiments with NAC, 1 mM was added for the final 2 days of differentiation.

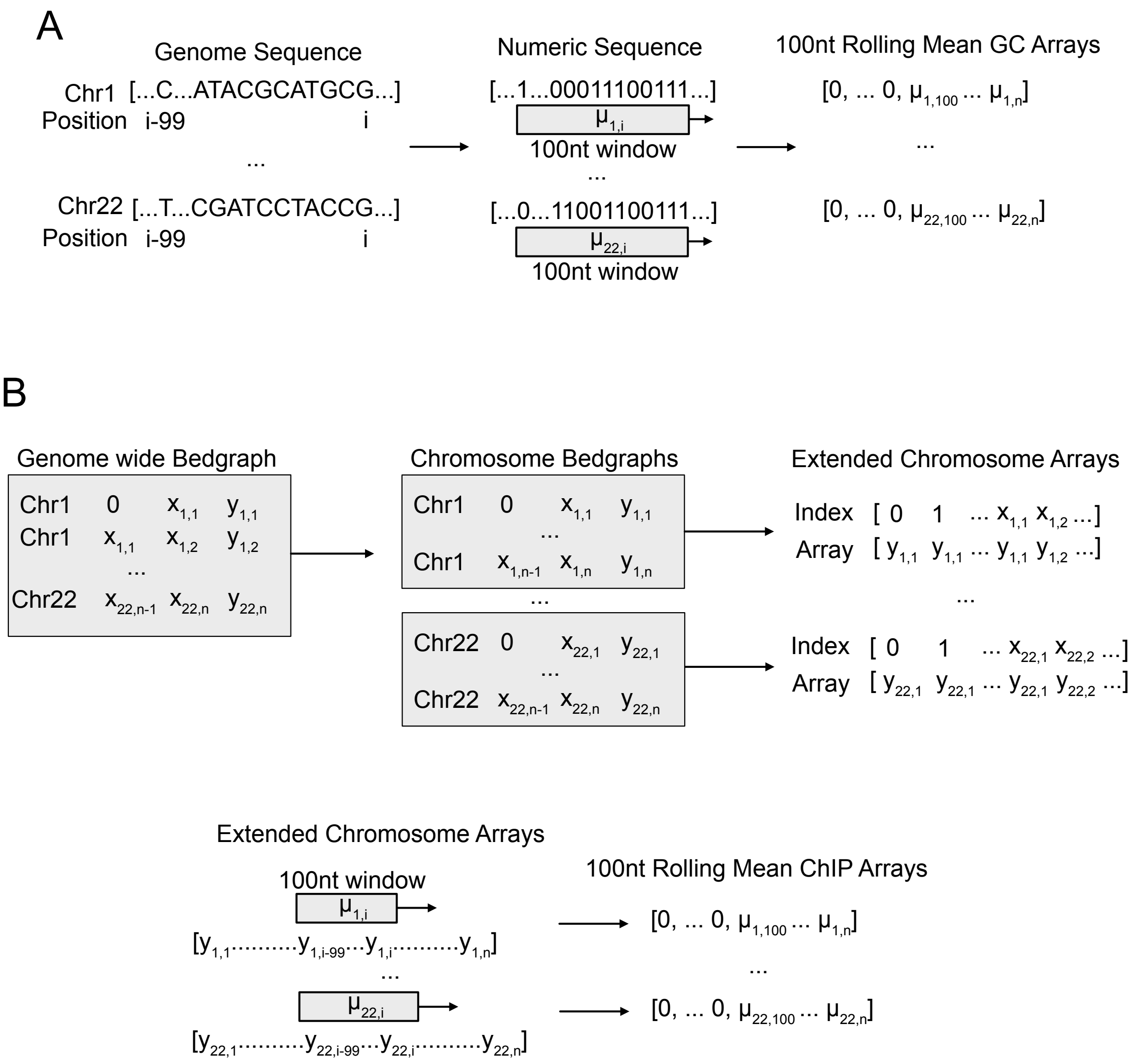


**Figure S10. Rolling mean ChIP strategy.** (A) Scheme to compute rolling mean GC content. Each chromosome sequence from GRCh38 was converted into a binary array with 0 for A/T and 1 for G/C. A 100nt window was used to compute the local GC content for each nucleotide of the genome. Due to the rolling mean algorithm, the first 99 (window size-1) nucleotides per chromosome were replaced with 0. (B) Scheme to compute rolling mean ChIP signal. The genome wide bedgraph output from MACS2 bdgcmp with the ppois flag was separated into chromosome bedgraph files. Bedgraphs were “extended” into Numpy arrays where each array index corresponds to a nucleotide position on the genome and the array value at each index is the bedgraph signal at that location. The 100nt rolling mean was computed for the extended array as in (A).

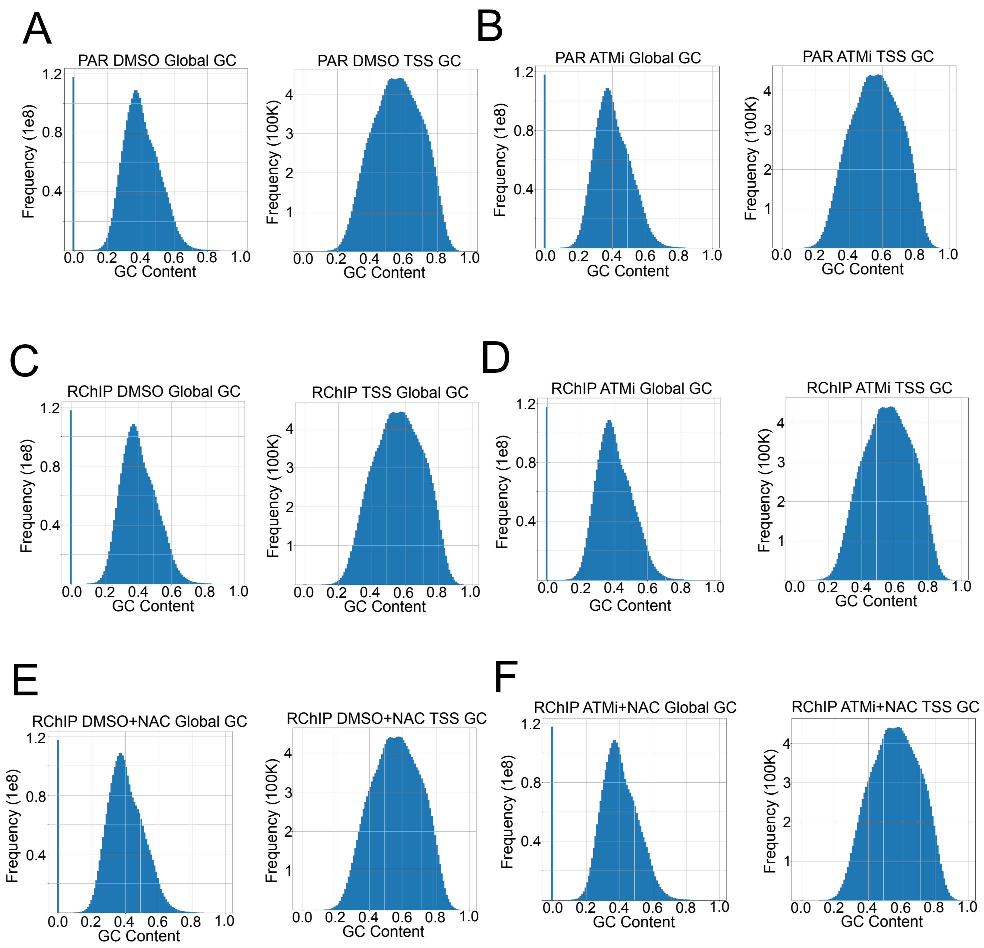


**Figure S11. Histograms for each subspace for each ChIP sample.** (A) Histograms for Global GC and TSS bins for DMSO PAR ChIP locations. (B) Histograms for Global GC and TSS bins for ATMi PAR ChIP locations. (C) Histograms for Global GC and TSS bins for DMSO RChIP locations. (D) Histograms for Global GC and TSS bins for ATMi RChIP locations. (E) Histograms for Global GC and TSS bins for DMSO+NAC RChIP locations. (F) Histograms for Global GC and TSS bins for ATMi+NAC RChIP locations.
